# Diet and Size-at-Birth Affect Larval Rockfish Condition and Survival

**DOI:** 10.1101/2024.04.07.588270

**Authors:** Kamran A. Walsh, Andrew R. Thompson, Garfield T. Kwan, Brice X. Semmens, H. William Fennie, Rasmus Swalethorp

**Affiliations:** Scripps Institution of Oceanography, University of California, San Diego, La Jolla, CA 92037; School for Marine Science and Technology, University of Massachusetts, Dartmouth, New Bedford, MA 02744-1221 (Present address); NOAA Fisheries Service, Southwest Fisheries Science Center, La Jolla, CA 92037-1508; University of California, Davis, Davis, CA 95616; NOAA Fisheries Service, Alaska Fisheries Science Center, Seattle, WA 98115

**Keywords:** ichthyoplankton, feeding ecology, prey selection, otolith growth, zooplankton

## Abstract

Feeding success and maternal effects on larval size have long been hypothesized as important contributors to interannual recruitment variability in marine fishes. This study examined the feeding ecology and influences of diet and size-at-birth on length and growth of larval rockfishes (*Sebastes* spp.). Prey carbon biomass and selection were calculated from gut contents, size-at-birth was estimated using otolith core size, and recent growth was derived from outer otolith increment widths. Biomass contributions of preferred prey and otolith data were integrated into Bayesian hierarchical models predicting length and growth. Larvae primarily fed on and selected for copepod nauplii and Calanoid copepodites, modulating feeding with ontogeny and in response to prey availability. Based on carbon weight, the relative contribution of Calanoid copepodites to the diet was more strongly and positively correlated with length and growth than that of nauplii. Younger larvae experienced faster growth in association with Calanoid copepodite consumption than older larvae. Positive effects of core radius suggest that initial larval size, believed to be mediated by maternal provisioning, increases the likelihood of survival, larger size and faster growth. These findings ultimately provide evidence that selective feeding and size-at-birth mediate rockfish survival in early life stages.

## 1. INTRODUCTION

While survival of marine fishes through the larval stage of development is generally thought to be important in driving adult populations, the specific factors that drive larval mortality remain largely unresolved (Lasker 1981, Robert et al. 2014, Hare 2014). Understanding the roles of different environmental factors in mediating larval growth, body condition and survival is therefore a critical part of efforts to connect early life history to fisheries recruitment and population dynamics. The capacity of larvae to find and feed on preferred prey has long been scrutinized. For instance, the seminal “critical period” (Hjort 1914, Hjort 1927) and “match-mismatch” hypotheses (Cushing 1990) propose that the strength of the spatio-temporal match between larvae and optimal prey determines year-class success. Next, the “growth-mortality” hypothesis posits larval recruitment variability within the context of surviving starvation and predation through fast growth and early recruitment (Anderson 1988). The complementary “bigger is better” and “stage duration” hypotheses propose that larger, faster growing larvae are more likely to survive (Chambers & Leggett 1987, Houde 1987, Meekan & Fortier 1996, Meeken et al. 2006). Slower-growing larvae are smaller for extended periods of time, and thus remain vulnerable to a wider range of ichthyoplankton predators than faster-growing larvae (Shepherd & Cushing 1980, Miller et al. 1988, Margulies 1989). The “growth-selective predation” hypothesis suggests slow growing larvae exhibit decreased escape responses when encountering predators due to the inherent physiological drawbacks of being in poor condition (Takasuka et al. 2003, Takasuka et al. 2007, Houde 2008). As a consequence, larger, faster growing larvae have a selective advantage over slower growing individuals (Hare & Cowen 1997). However, conclusive *in situ* evidence supporting the above hypotheses in relation to prey availability remains limited (Leggett & Dublois 1994). Moreover, studies have failed to reveal positive relationships between bulk zooplankton biomass and recruitment success (e.g. Agostini et al. 2007, Irigoien et al. 2009). As such, more studies are needed to investigate whether larval diet selectivity is indeed a driver of recruitment (Robert et al. 2014).

Diet selectivity is manifested as tradeoffs between energetic input gained from prey consumption and energetic output spent on searching and prey capture. In general, the few studies that found positive relationships between prey availability and indices of recruitment strength or larval body condition also resolved prey taxonomy (e.g., Beaugrand et al. 2013, Murphy et al. 2012, Malca et al. 2022). However, the prey types most conducive to growth and survival are not always the most strongly selected by larval fishes (Burns et al. 2021). Moreover, low abundances of preferred prey may result in a switch to another taxa (Anderson 1994). This is further complicated by shifts in zooplankton community composition and timing in response to changing oceanographic conditions, and may result in changes in diet selection with potential negative consequences to growth and condition (Anderson 1994). Despite the clear importance of evaluating larval feeding ecology and its impact on recruitment success, a conclusive understanding of prey selection’s influence on larval growth and survival remains elusive.

Maternal provisioning is another factor believed to affect recruitment success, with observed positive effects on larval size-at-birth, endogenous energy stores, body condition and survival (Hislop 1988, Chambers & Leggett 1996, Berkeley et al. 2004, Fennie et al. 2023).

Otolith core size (aka nuclear width, hatch check diameter) has been correlated with size-at-birth and early life survival in multiple species (Meekan & Fortier, 1996, Garrido et al. 2015, Fennie et al. 2023), and this metric has been recently proposed to be a robust indicator of maternal provisioning. Through this, we can use otolith core size to assess the effects of size-at-birth on larval condition and performance. To the best of our knowledge, there are few laboratory studies that simultaneously examine the impacts of maternal provisioning and early life diet on larval condition (Perez & Fuiman 2015, Kolodzey et al. 2021), and none that use otolith core size as a metric of maternal effects or examine these relationships in wild-caught larvae.

Rockfishes are a genus of ovoviviparous Scorpaeniformes abundant in North Pacific waters and throughout the California Current Ecosystem (CCE). The genus *Sebastes* is diverse in morphology, size, habitat, and behavior, ranging from larger economically valuable species heavily targeted in commercial and recreational fisheries to smaller counterparts that are either not subject to fishing or are common as bycatch. Rockfishes of all life stages are also important prey for higher trophic level predators (Love et al. 2002, Field et al. 2007). Due to the long lifespans, relatively low fecundity, and slow growth of many rockfishes, they are highly vulnerable to overexploitation and stocks of several species suffered dramatic declines over the past century (Love et al. 1998, Butler et al 2003). However, many responded positively to conservation efforts including spatial closures (Thompson et al. 2017, Keller et al. 2019, Freeman et al. 2022). In addition, two recent studies provided important insight into conditions that facilitate rockfish recruitment. First, Schroeder et al (2019) found that recruitment for multiple rockfish species correlated with the exposure of adults to cold, high oxygen, high nutrient, low salinity Pacific subarctic water (California Current water) in the months prior to recruitment. Second, Fennie et al (2023) demonstrated that females spawn larger larvae when bathed in Pacific subarctic water during gestation and that larvae born larger were more likely to survive the first few weeks of life. We build on these findings to evaluate the combined role of size-at-birth and larval prey on growth and condition. Larval *Sebastes* are known to prey on copepod eggs, nauplii, and copepodites, as well as euphausiids, diatoms, and protists (Sumida & Moser 1984, Burns et al. 2021). However, the degree to which larvae in the CCE select for these prey items, whether preference involves the active selection of specific prey taxa or passively selecting organisms based on their relative abundances in the surrounding environment, and the explanatory power of both prey consumption and otolith core size on their size and growth is unknown.

Here, we investigate the impacts of development stage-specific prey selectivity and size-at-birth on larval size and recent growth. The primary objectives of this study are to (i) describe the diet of *Sebastes* larval assemblages within the Southern California Bight, (ii) examine ontogenetic trends of niche breadth, prey preference, and active/passive selection, and (iii) determine the degree to which size-at-birth and the consumption of different prey taxa contribute to size, growth, and survival at multiple ages.

## 2. MATERIALS AND METHODS

### 2.1 Sample Collection

Larval rockfishes were collected along seven inshore California Cooperative Oceanic Fisheries Investigations (CalCOFI) sampling locations (Line-Station 85-42.9, 86.7-33, 90-30, 90-35, 90-37, 93.3-28, and 93.3-35) in the Southern California Bight in 2021 (Figure 1). Stations 93.3-28 and 90-30 were sampled in both winter and spring, station 86.7-33 was sampled only in winter, and all other stations were sampled only in spring. Samples were collected via oblique tows from the upper 30 m of the water column using a 70-cm bongo (bongo-70) frame with 505- µm synthetic nylon mesh nets (Smith & Richardson 1977). The volume filtered was determined using a mechanical flowmeter centered in the mouth of the starboard net. Upon collection, the cod ends of the bongo nets were immersed in a saltwater ice slurry for one minute to hopefully increase stomach content retention of the captured larvae. Following this, the samples were immediately concentrated using cold, unfiltered seawater through a 300µm sieve and fixed in 95% tris-buffered ethanol (EtOH). To characterize the prey field available at each sample station, a second bongo with 202-µm mesh and 53-µm mesh nets was deployed shortly after each 505 µm haul at stations 90-35, 90-37, 93.3-35, and 93.3-28 (both winter and spring). A second flowmeter was centered in the mouth of each bongo. The contents of both nets were filtered through sieves, and samples from each net were fixed in 3.7% formalin to ease identification of small zooplankton.

**Figure 1:**
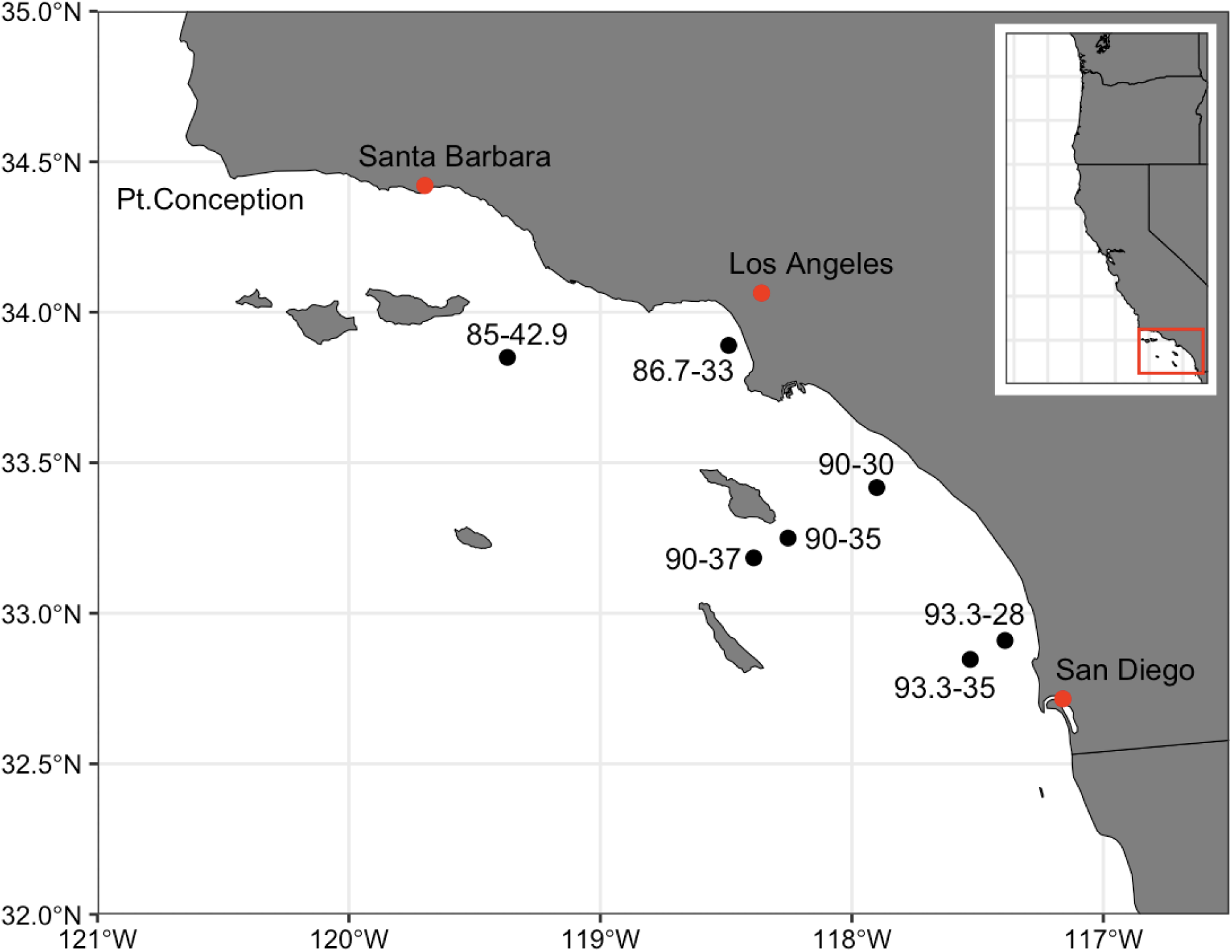
Schematic of California Cooperative Fisheries Research Investigations (CalCOFI) sample stations of ichthyoplankton and prey field collection in the Southern California Bight (SCB).

### 2.2 Larval Rockfish Identification

Rockfish larvae were sorted from the 505-µm samples under a dissecting microscope, photographed, and identified to either preflexion (after yolk sac absorption and before start of notochord flexion), flexion (start of notochord flexion to notochord tip angle ∼ 45◦) or postflexion (completion of notochord flexion) stages. Following this, standard length (SL) was measured to the nearest 0.01mm from the tip of the lower jaw to the tip of the notochord in preflexion and flexion larvae and to the distal-most hypural margin in postflexion larvae using an eyepiece micrometer.

Because most rockfish larvae are indistinguishable to species based on morphology (Moser 1996), they were genetically identified via Sanger Sequencing as described in Thompson et al. (2016). Briefly, tissue samples were taken from the eye of the larvae, and chelex-based boiling was used to extract genomic DNA from each sample (Hyde et al. 2005). The target genomic region was amplified by polymerase chain reaction (PCR) using the primers GLURF2- 5’ and CB3RF2-5’ (Hyde et al. 2008). Each PCR was conducted in 10µl volumes with buffer (67mM Tris-HCl pH 8.8, 16.6mM [NH4]2SO4, 10mM β-mercaptoethanol, 2mM MgCl2), 800µM dNTP, 0.4µM of each primer, 0.5mg ml^−1^ bovine serum albumin, 0.5 units Taq DNA polymerase (New England Biolabs), and 1µl of chelex supernatant containing DNA template. The thermal profiles of the PCRs were: denature at 92°C for 2 min and 30 sec; followed by 40 cycles of 94°C for 30 sec, 55°C for 90 sec, 70°C for 90 sec; then a final extension of 72°C for 3 min. Negative, no template controls were run for each PCR to monitor for possible contamination. PCR products were enzymatically cleaned using ExoSap-IT (Affymetrix). The resultant products were sequenced using the internal primer CBINR3 (5’-ATG AGA ART AGG GGT GGA AGC T-3’) and BigDye v.3.1 Dye Terminator chemistry following manufacturer’s protocols. The sequenced products were analyzed using an ABI3730 Genetic Analyzer (Life Technologies). Finally, the sequences were edited and aligned using Sequencher v.4.9 (GeneCodes), aligned with templates from reference adult rockfishes (Hyde & Vetter 2007), and identified by creating Neighbor Joining phylogenetic trees with MEGA v11 (Tamura et al.

2021).

### 2.3 Stomach Content Analysis

The entire digestive tracts of 161 larvae were removed and dissected. The gut contents were identified, photographed, and lengths and widths measured to the nearest 0.001mm using an eyepiece micrometer. All recovered prey items were identified to the taxonomic class or order level because their partially digested condition made it difficult to reliably determine genus or species. Those that could not be reliably identified to a class or order level were not included in subsequent analyses. To measure relative carbon (C) biomass contributions of respective prey taxa, carbon biomass weights in µg C were estimated for each recovered prey item using length- dry weight conversion factors from existing literature (Supplementary Table 1). The carbon biomass consumed by each larva was used as a proxy for feeding success. Recovered prey items for which accurate length and width measurements were not possible were not assigned carbon biomass values or used in size range feeding selectivity calculations.

### 2.4 Zooplankton Identification and Enumeration

Plankton community samples from the 202-µm mesh nets were analyzed via ZooScan digital scanning and image analysis (Gorsky et al. 2010). Subsamples were taken for each jar and were split into small (202-999 µm), large (1000-4999 µm), and extra-large (5000-999999 µm) sizes using a 1-mm and 5-mm mesh sieve. Two aliquots of the small size range, one aliquot of the large size range, and one aliquot of the extra-large size range were scanned at 2400 dpi resolution (four scans per sample). In total, 29,165 vignettes were classified by a machine learning algorithm using Random Forest classification and regression tree methods, and all images were reexamined manually to correct misidentifications. Ten individuals of each prey taxon identified from the gut contents of larval *Sebastes* spp. from each subsample size at each station were randomly selected and body length (prosome length for copepodites) and width measured to the nearest 0.001µm using ImageJ image processing software.

Plankton samples from the 53-µm mesh nets were identified and enumerated manually under a dissecting microscope due to limitations in ZooScan optical resolution. Aliquots were removed by Hensen stempel pipette, analyzed and organismal counts standardized to number per m^3^. Approximately 30 individuals of each commonly consumed prey taxon and approximately 10 individuals of all other taxa from each station were randomly selected for size measurement using an eyepiece micrometer. All plankton identifications in ZooScan and manual microscopy processed samples were made using regional taxonomic literature, with taxa identified to the class or order level consistent with the taxonomic resolution of the stomach contents. *In situ* zooplankton C weights from both the 202-µm (ZooScan) and 53-µm (microscopy) samples were estimated using the same length-weight conversion factors as previously described (Supplementary Table 1). Taxon-specific abundance in counts per m^3^ and C biomass in µg C per m^3^ were calculated for the four sample stations in which larval feeding selectivity was calculated, both to compare with gut contents and to gain a sense of spatial and temporal plankton community variation.

Data from the 202-µm and 53-µm samples were compared to determine size cutoffs based on the respective capture efficiencies of each mesh size for assessing plankton abundance within different length and width ranges. Size cutoffs were made by comparing the abundances of each taxon and developmental stage at a given length or width range in the 202-µm samples to the abundances of the same taxonomic group and size range in the 53-µm samples (Table 2).

Taxa categories of a given size range with greater abundances in one mesh size than the other were used for subsequent analyses, with length and width range cutoffs used for the length and width range selection calculations, respectively.

### 2.5 Otolith Analysis

Sagittal otoliths from preflexion, flexion and postflexion larvae (n = 80) were extracted under a dissecting microscope and mounted on a microscope slide. Individuals were selected haphazardly to represent all seven sample stations. Otoliths were photographed using a microscope with an oil immersion lens at 100x magnification and rendered via HeliconFocus v8.1.2. Due to a lack of substantial evidence indicating variation in otolith microstructure between left and right sagittae, the otolith image with the clearest resolution from each larva was selected for further analysis. Otolith microstructure was measured using the RFishBC package in R (Ogle 2022). Core radius was measured from the center of the otolith outwards to the first visible band, age was measured by the number of daily increments from the first visible band to the outer edge of the otolith, and recent growth was estimated by the average widths of the three outermost complete daily increments-at-age. Multiple reads were taken by two readers and compared. Additional reads were taken as needed to determine final ages for use in subsequent analyses.

### 2.6 Data Analysis - Diet, Feeding Niche, Prey Preferences

We first evaluated whether there were stage-specific differences in diet. Differences in average gut content carbon biomass of prey (C biomass ∼ development stage), number of prey (no. prey ∼ development stage), and average size of prey (prey size ∼ development stage) between developmental stages were assessed via one-way analysis of variance (ANOVA) in R version 4.3.2. Relative carbon contributions of prey were then calculated for all individual larvae in each growth stage and at each sample station based on the eleven most common prey groups identified: Calanoida nauplii, Cyclopoida nauplii, other Copepoda nauplii, Cladocerans (Podonidae), Calanoida copepodites, Cyclopoida copepodites, Poecilostomatoida copepodites, other Copepoda copepodites, Copepod eggs, Euphausiid nauplii and Euphausiid juveniles (calyptopsis and furcilia stages). For the initial diet composition analyses, ‘other Copepoda Nauplii’ and ‘other Copepoda Copepodites’ included Eucalanid nauplii, Harpacticoids, and copepods not identifiable to a higher taxonomic resolution. Differences in gut content C biomass in each taxonomic category between growth stages, stations, and rockfish species were calculated using permutational multivariate ANOVA (PERMANOVA) in Primer v6.1.7 (Primer, LTD) on a Bray-Curtis similarity matrix with 999 permutations. Growth stages were nested within stations and species identity was treated as a random effect to assess spatial and/or species variation in diet for each growth stage. PERMANOVA tests were run both with all *Sebastes* species pooled and with individual species included as a random factor. To evaluate which prey taxa drove significance in the PERMANOVA, similarity percentage (SIMPER) analysis was performed.

To evaluate if rockfish larvae targeted specific prey at different life stages, we calculated three indices of prey choice. Dietary indices, taxonomic niche breadth, and feeding selectivity were evaluated using the Index of Relative Importance (%IRI), Levins’ standardized niche breadth, and Chesson’s selectivity index (α), respectively. Dietary indices and niche breadth were calculated for each larval developmental stage based on eleven prey categories. For Chesson’s selectivity index, the prey categories were reduced to the seven most commonly observed taxa. Unidentified Copepoda nauplii were removed in assessing prey preference.

Cladocerans were removed due to insufficient *in situ* abundances from three of the four stations analyzed for selectivity. Euphausiid nauplii and post-naupliar stages were grouped together, and Poecilostomatoids were included in the ‘Other Copepodites’ category.

To assess the degree to which major contributors of total prey abundance and biomass were consumed throughout the sample population, an Index of Relative Importance was estimated as:

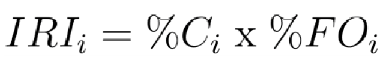

in which (%𝐶_𝑖_) is the percentage contribution of each prey category *i*’s contribution to the total abundance or biomass consumed by each larval growth stage, and (%𝐹𝑂_𝑖_ ) is the percentage of larvae with non-empty guts in a given growth stage that had ingested that prey category. 𝐼𝑅𝐼_𝑖_was calculated for all prey categories *i* and the relative importance *%*𝐼𝑅𝐼_𝑖_ of each prey category was calculated as:

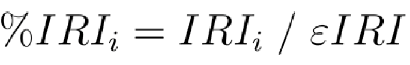

where ε𝐼𝑅𝐼_𝑖_ is the sum of all 𝐼𝑅𝐼_𝑖_ estimates.

To assess if rockfish feeding became more generalized or specialized with development, Levins’ standard niche breadth was calculated as:

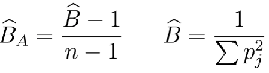

Where. 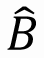 reflects niche breadth, 𝑝_𝑗_ is the proportion of the diet consisting of prey category *j,* and *n* is the total number of prey categories observed. High. 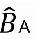 values are an indication of a wide taxonomic feeding niche, indicating generalist feeding behavior, while low values reflect a narrow niche and specialist feeding behavior.

Selectivity calculations were based on the sample stations in which the *in situ* samples had been collected to describe prey preference across ontogeny, across spatial scales, and with respect to size ranges of consumed prey. Selectivity of all individuals from all stations was pooled to gain a sense of trends in prey preference, selectivity within each sample station was calculated to compare spatial differences in feeding preference, and selectivity within prey size ranges was calculated using logarithmic size intervals, consisting of eleven length classes (length midpoint (µm) - 84, 119, 168, 237, 335, 473, 668, 944, 1334, 1884, 2661, corresponding length ranges - 72-100, 101-141, 142-200, 201-282, 283-400, 401-562, 563-800, 801-1122, 1123-1585, 1586-2239, 2240-3162) and nine width classes (width midpoint (µm) - 60, 84, 119, 168, 237, 335, 473, 668, 944, corresponding width ranges - <71, 72-100, 101-141, 142-200, 201-282, 283- 400, 401-562, 563-800, 801-1122) using measurements from the ZooScan and manual microscopy. To estimate feeding preference, Chesson’s α-selectivity index was calculated as (Chesson 1978):

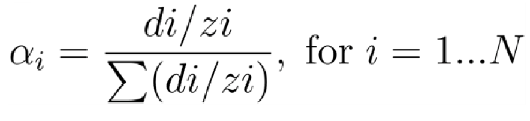

where *di* is the abundance of prey type *i* in the guts, *zi* is the abundance of that prey item *i* in the environment, 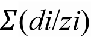 is the ratio of gut content to *in situ* abundance for all prey categories, and *N* refers to the number of prey categories *i*. A threshold of 14.3% ((1/prey categories *N*)*100) was estimated as neutral preference for measures of taxa selection, with smaller values indicating negative (weak) selection and larger values indicating positive (strong) selection. For sample station selection calculations, a threshold of (1/[prey categories *Np* x number of stations *Ns*]*100) was estimated as neutral preference. For size range categories, a threshold of (1/[prey categories *Np* x prey size categories *Ns*]*100) was used. Values of α*i* were calculated for each individual larva and averaged by developmental stage. PERMANOVA and SIMPER analyses were used to compare prey selection between growth stages, stations and species. Similarly as with gut content C biomass, growth stages were nested within station and separate PERMANOVA tests were run with all *Sebastes* spp. grouped together and with individual species as a random factor.

Due to the small sample sizes of many species, it was necessary to lump all individuals into a single *Sebastes* spp. group for all analyses (IRI, Levins’, Chesson’s, ANOVA) except for the PERMANOVA diet and selection tests. By pooling species, we made the assumption that different species have similar diets, co-occur in space and time and are thereby exposed to similar prey communities and levels of maternal investment which is partly supported by past studies (Bosley et al. 2014, Thompson et al. 2016, Ralston et al. 2013, Schroeder et al. 2019). Moreover, PERMANOVA results (see Results section) showed no species effect.

### 2.7 Data Analysis – Influence of Diet and Otolith Core Size on Larval Condition

A Bayesian hierarchical model was used to evaluate the effect of otolith core size on the age of fish, a proxy for survival (model 1). Diet and otolith data were integrated into two additional hierarchical models to explore the relative effects of age, the relative contributions of different prey taxa in the diet, and size-at-birth on model 2) larval size (standard length) and model 3) growth (averaged width of the outer 3 otolith bands). Four Hamiltonian Markov chains (MCMC) with 1000 iterations each were used. We standardized all dependent and independent variables, and thus a mean of 0 and standard deviation of 1 were initially included as informative priors. Normally distributed posterior distributions were summarized using means and standard deviation. The data were fit to the following equations:

Age ∼ Size-at-Birth Model (1):

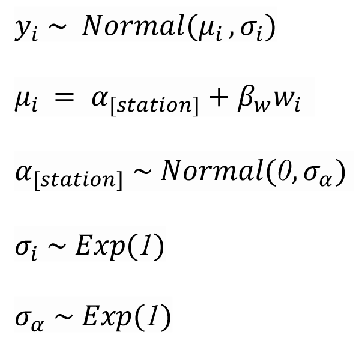

Integrated Diet, Size-at-Birth, and Age Models (2 & 3):

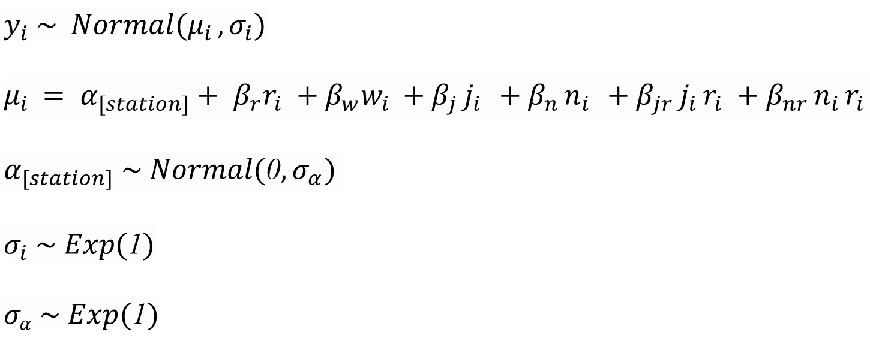

In all models, sample station number was entered as a random effect to account for any spatial differences in environmental conditions, size-at-birth, and prey field biomass or community composition (Table 1). Core radius was entered as a fixed effect in model 1. Core radius, the ratio of Calanoid Copepodite C biomass over total C biomass, and the ratio of Calanoid and Cyclopoid Nauplii C biomass over total C biomass were entered as fixed effects in models 2 and 3. All fixed effects were standardized (mean 0, standard deviation 1). Proportions of prey categories over the total C biomass consumed by each fish were used to determine how consumption of certain prey groups relative to the total biomass of consumed prey affected size and growth. Interactions between age and each diet parameter were integrated into the models to account for expected increases in C biomass uptake with age, and to determine how diet affects condition at different stages of larval development. Interactions between otolith core and age were not included as maternal effects are not affected by the age of the progeny. Predictive accuracy of the models to the observed data was determined using posterior predictive distributions, and MCMC convergence diagnostics were graphically computed using traceplots.

**Table 1:** parameters used in modeling framework.

| Parameter | Description | Units |
| --- | --- | --- |
| $y$ | Age (model 1), Standard Length (model 2), Recent Growth (model 3) | Days, mm, $\mu\text{m}$ |
| $\alpha$ | Intercept | |
| $station$ | Sample station | Categorical integer |
| $r$ | Age (models 2 and 3) | Days |
| $w$ | Otolith core radius, radius from center to extrusion check | $\mu\text{m}$ |
| $j$ | Proportion of Calanoid copepodite gut content carbon biomass over total carbon biomass | % $\mu\text{g C}$ |
| $n$ | Proportion of Calanoid and Cyclopoid nauplii gut content carbon biomass over total carbon biomass | % $\mu\text{g C}$ |

**Table 2:** Size range cutoffs for length and width measurements from 53µm and 200µm nets. ‘53µm max’ represents the maximum size range in which abundances of each taxon in the 53µm net were higher than in the 200µm, while ‘200µm min’ represents the minimum size range in which abundances in the 200µm net were higher than in the 53µm.

| Length |  |  | Width |  |  |
| --- | --- | --- | --- | --- | --- |
| Prey Taxa | 53µm max. size | 200µm min. size | Prey Taxa | 53µm max. size | 200µm min. size |
| All copepodites | 562 | 563 | All copepodites | 200 | 201 |
| All nauplii | 282 | 283 | All nauplii | 200, 401-562 | 201-400 |
| Euphausiid juveniles | - | All | Euphausiid juveniles | - | All |
| Cladocerans | - | All | Cladocerans | - | All |
| Eggs | 200 | 201 | Eggs | 200 | 201 |

The modeling framework was coded in the R ‘rethinking’ package (McElreath 2020). All data used for this project are posted on BCO-DMO https://www.bco-dmo.org/project/867657, and code is available at https://github.com/KamranWalsh/SCB-larvalrockfish.

## 3. RESULTS

### 3.1 Rockfish Assemblage Species and Size Structure

Analysis of gut content was performed on 161 intact larval rockfish, encompassing 21 species representing a range of sizes, morphologies, and habitat types (Supplementary Table 2). The most abundant species were *Sebastes semicinctus* (Halfbanded Rockfish), *S. hopkinski* (Squarespot Rockfish), and *S. jordani* (Shortbelly Rockfish). The most frequently occurring species were *S. semicinctus* and *S. simulator* (Pinkrose Rockfish). The sample set consisted of 55 preflexion stage (mean SL = 4.59 mm ± 0.78 standard deviation unless otherwise noted), 61 flexion stage (mean SL = 6.51 mm ± 1.27), and 45 postflexion stage (mean SL = 9.47 mm ± 1.92) larvae. Size ranges between preflexion (3.03-6.49 mm SL), flexion (3.99-9.23 mm SL) and postflexion (6.42-15.42 mm SL) stages overlapped. While larvae were selected for gut content analysis in an effort to provide uniformity in ontogenetic distributions representative of variability between sample stations (Supplementary Table 3), this was constrained by uneven spatial distribution of growth stages.

### 3.2 Larval Diet

In total 3,930 prey items were recovered from the 161 larvae. Of these, 3,074 were sufficiently intact to allow identification and use in subsequent analyses. Only 7 individuals had empty guts: 6 preflexion larvae (SL = 3.54-5.68 mm) and one flexion larva (SL = 5.76 mm).

There were significant differences in the total number (df = 2, F = 53.50, p<0.001), average size (df = 2, F = 53.95, p<0.001) and average C biomass (df = 2, F = 12.47, p<0.001) of prey consumed in different stages of ontogeny. As larvae grew, their gut contents included prey of increasing size and diminishing contributions of smaller sizes (preflexion = 207.2 µm ± 61.6, flexion = 247.5 µm ± 107.6, postflexion = 395.2 µm ± 255.0), increasing quantity (preflexion = 5.00 ± 4.06, flexion = 19.36 ± 13.61, postflexion = 35.96 ± 22.87) and greater average C biomass (preflexion = 0.175 µg ± 0.200, flexion = 0.259 µg ± 0.557, postflexion = 1.23 µg ± 4.75), with an overall 7.2x increase in number of prey items contained in the gut between early preflexion and late postflexion stages.

The three most commonly consumed prey taxa across all stages of larval development were Calanoida nauplii, Cyclopoida nauplii, and Calanoida copepodites, which accounted for 83.5% of prey items and 70.2% of the total C biomass. Calanoida nauplii were the single most common prey type (32.9%), and Calanoida copepodites comprised the largest portion of gut content C biomass (59.4%). The remainder of the identifiable gut contents consisted primarily of other copepodites and nauplii, copepod eggs, Euphausiid naupliar, calyptopsis, and furcilia stages, Cladocerans, Tintinnids and larval Bivalves.

Carbon biomass contributions of prey taxa differed significantly between sample stations (df = 8, F = 3.33, p = 0.001). Dissimilarities in C biomass contributions of prey groups between stations were primarily driven by Calanoid copepodite C biomass. Taxa also responsible for driving spatial C biomass dissimilarities were Calanoid nauplii and copepod eggs. Prey C biomass differed significantly between larval developmental stages nested within sample stations (df = 15, F = 3.95, p = 0.001), while the degrees of dissimilarity between preflexion/flexion (69.90%) and flexion/postflexion (67.32%) larval C biomass taxa contributions were similar.

Preflexion to flexion shifts were characterized by increased consumption of Calanoid nauplii relative to other prey types, which accounted for 19.91% average dissimilarity in gut content C biomass. Flexion to postflexion shifts were characterized by increased consumption of Calanoid copepodites, accounting for 26.94% average dissimilarity between the two growth stages. The prevalence of Calanoid copepodites in the diet increased with development. Calanoid copepodites were the dominant prey in postflexion larvae and represented the highest proportion of C biomass in both flexion and postflexion larvae (Figure 2a,c). Euphausiid calyptopsis and furcilia stages were also in the gut contents of some of the larger larvae, contributing to much of the C biomass unaccounted for by Calanoids in postflexion larvae. When taking %IRI dietary indices into account, the importance of nauplii diminished with ontogeny while the relative contribution of Calanoid Copepodites to the diet generally increased. The low contribution of Euphausiids to the %IRI indicates that relatively few larvae fed on Euphausiids, despite the high energy content of these prey (Figure 2b,d). No significant differences in C biomass contributions of prey were found between different species of rockfish.

**Figure 2:**
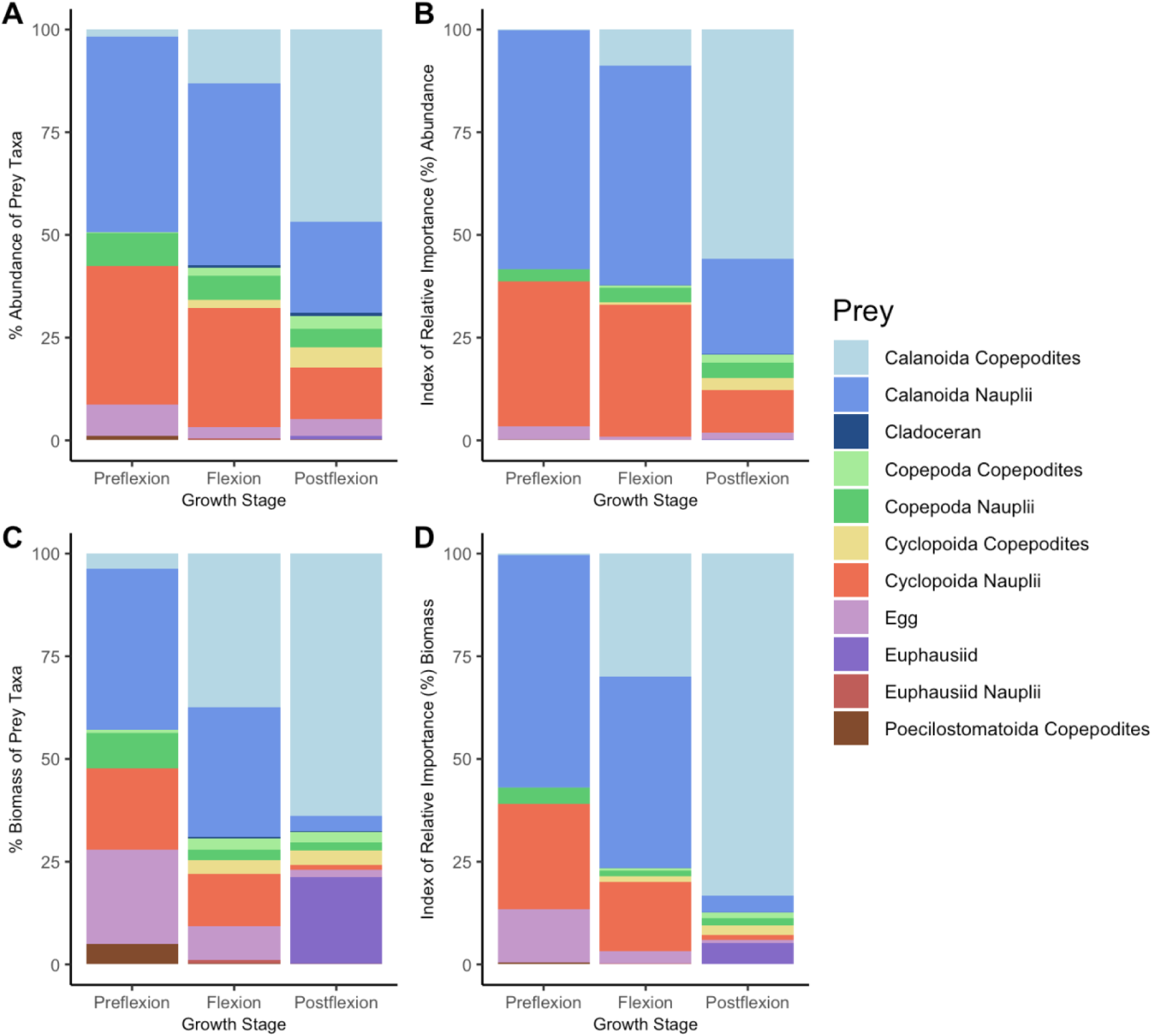
Proportions and Index of Relative Importance (%IRI) of the most commonly consumed prey groups within the gut contents by abundance (A, B) and gut content carbon biomass (C, D).

### 3.3 Taxonomic Niche Breadth and Prey Preferences

Feeding niche generally narrowed with increased development, with notable exceptions between flexion and postflexion larvae at station 90-37 and preflexion and flexion larvae at station 90-35 (Figure 3). While flexion and postflexion larvae often fed on a larger number of prey types than their respective preceding developmental stages (Figure 2), a larger proportion of their diet generally consisted of fewer prey taxa than observed earlier in ontogeny. However, the degree to which this occurred varied spatially.

**Figure 3:**
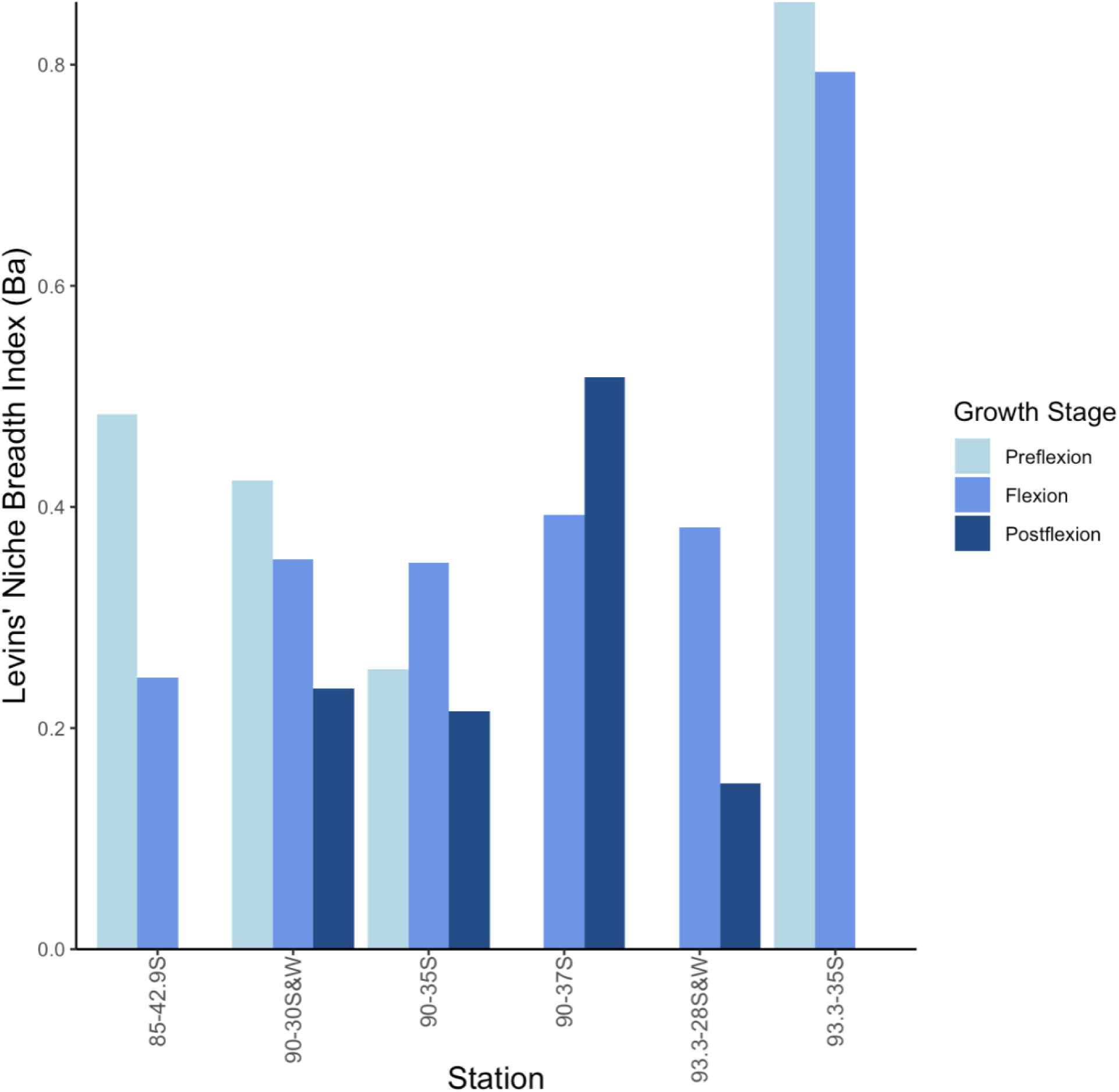
Levins’ standardized niche breadth (BA) illustrating changes in feeding niche between developmental stages across six sample stations. Growth stages of a particular sample stage with n<3 were excluded.

Taxonomic feeding selectivity also changed with ontogeny (Figure 4) and in response to prey availability in the surrounding environment. There were significant differences in preference across sample stations (df = 4, F = 4.78, p = 0.001) and between growth stages nested within stations (df = 9, F = 2.42, p = 0.003), indicating modulation of selection in response to changing abundances of prey taxa in ambient zooplankton communities. Significant differences in preference between flexion and postflexion larvae were observed in three of the stations (90-37: p = 0.005, 93.3-35: p = 0.001, 90-35: p = 0.049), but no significant differences between preflexion and flexion larvae were observed in any stations. There were no detectable differences in preference between species. The general trend in diet was driven by a strong selection towards copepod eggs and nauplii in first feeding larvae and a gradual transition in selectivity towards Calanoid copepodites with larval ontogeny (Figure 4). Greater dissimilarity in preference between flexion and postflexion larvae (63.54%) was observed than between preflexion and flexion larvae (50.25%). Selection towards Calanoid nauplii was strong across all sample stations and growth stages, while selection towards Cyclopoid nauplii was more prevalent in preflexion and flexion larvae. Although larvae did not select for Cyclopoid copepodites, other copepodites, and Euphausiids on a sample population level, these prey groups were still selected for in some sample stations (Table 3).

**Figure 4:**
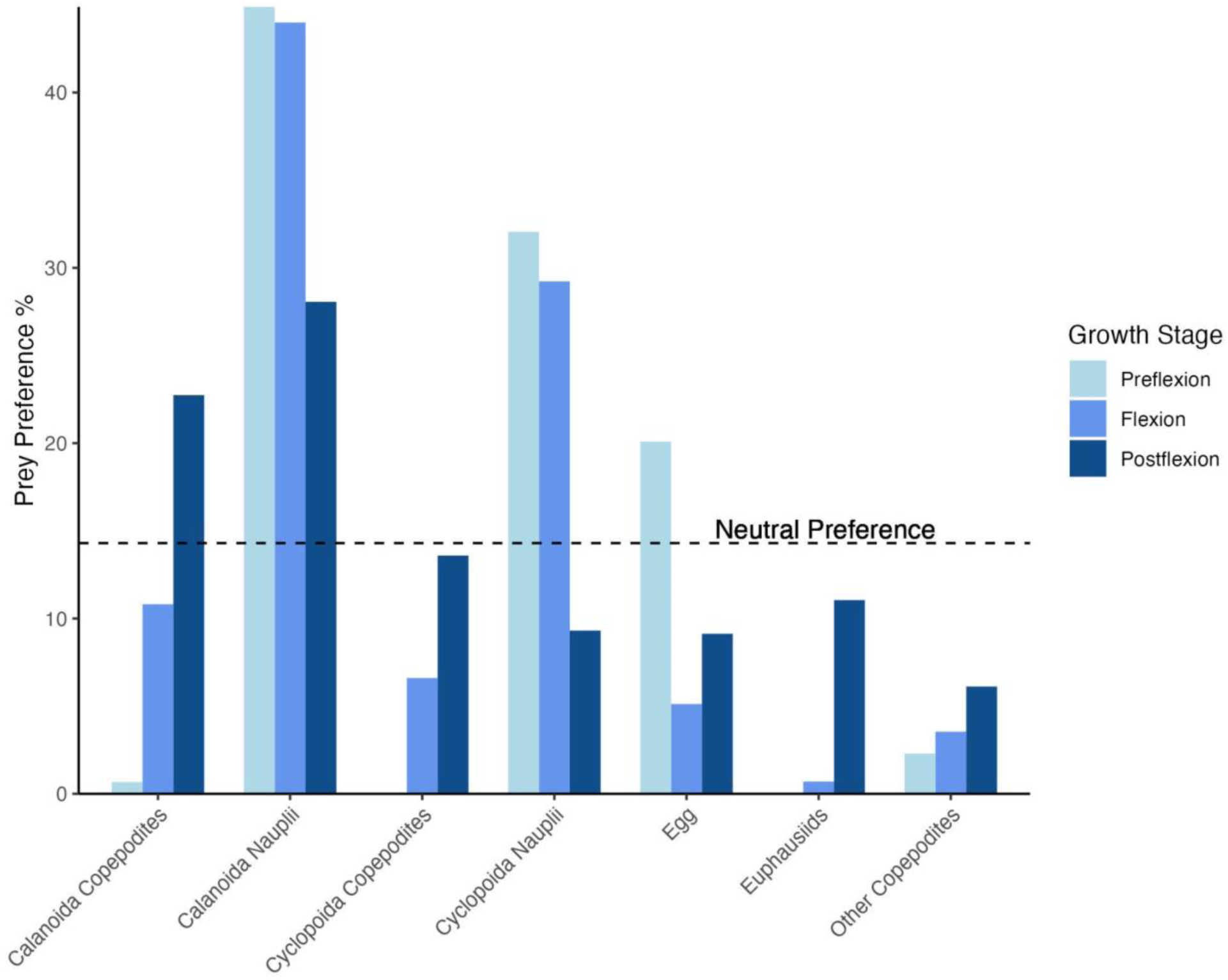
Prey taxonomic preferences of larval rockfish development stages. The dashed line indicates the neutral prey preference % threshold of 14.3%.

**Table 3:**
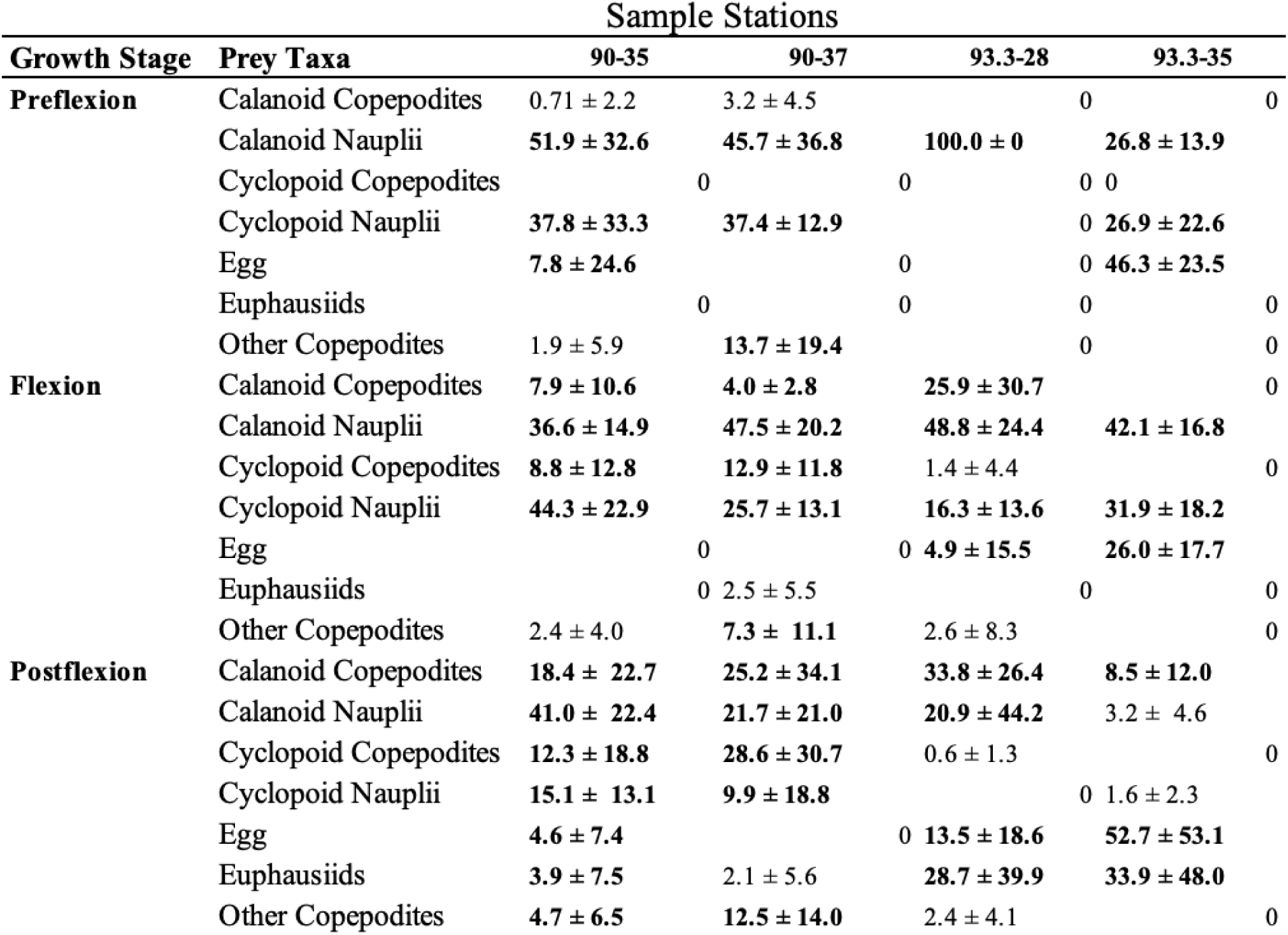
Chesson’s α-selectivity index calculations for four sample stations with the seven most commonly consumed prey taxa. Values are average α ± standard deviation. Bolded values represent taxa and size ranges that are positively selected for. Neutral preference is 3.57%.

Larvae also exhibited strong preferences towards specific size classes of their prey, but the degree to which selection was centered around specific length and width ranges differed between growth stages. When selection was calculated for each logarithmic length and width range (Tables 4 and 5), larvae generally exhibited stronger selection towards specific width classes and weaker selection across a wider range of length classes. While the preferred size ranges of prey increased with ontogeny, size ranges within the most strongly selected for prey groups did increase as dramatically as expected. Rather, selection generally followed copepod developmental history, with the strongest selection across different growth stages for given prey groups often falling within relatively similar size ranges. Larger larvae began to incorporate larger size ranges in their feeding, but nevertheless displayed considerable overlap in size range of prey with preceding growth stages. Width class selection for all three growth stages was generally strongest in the 142-200µm range, and length class selection was strongest primarily in the 201-282µm range. Pronounced size range increases with ontogeny also occurred as the result of the inclusion of different prey groups into the diet of older larvae, with postflexion larvae exhibiting stronger selection for Euphausiid nauplii and juveniles, Cyclopoid copepodites, and other copepodites in some size ranges.

**Table 4:** Chesson’s α-selectivity index calculations for eleven logarithmic length classes with the seven most commonly consumed prey taxa. Length class with midpoint 84µm had no values and was omitted from table. Values are average α ± standard deviation. Bolded values represent taxa and size ranges that are positively selected for. Neutral preference is 1.30%.

| | | Prey Length Midpoints (μm) | | | | | | | | | |
| --- | --- | --- | --- | --- | --- | --- | --- | --- | --- | --- | --- |
| Flexion Stage | Prey Taxa | 119 | 168 | 237 | 335 | 473 | 668 | 944 | 1334 | 1884 | 2661 |
| Preflexion | Calanoid Copepodites | 0 | 0 | 0 | 1.7 ± 7.3 | 0.1 ± 0.5 | 0 | 0 | 0 | 0 | 0 |
|  | Calanoid Nauplii | **2.2 ± 9.7** | **11.7 ± 24.5** | **24.1 ± 31.1** | **2.3 ± 10.3** | 0 | 0 | 0 | 0 | 0 | 0 |
|  | Cyclopoida Copepodites | 0 | 0 | 0 | 0 | 0 | 0 | 0 | 0 | 0 | 0 |
| | Cyclopoida Nauplii | 0.4 ± 1.9 | **5.3 ± 8.0** | **31.0 ± 34.7** | 0 | 0 | 0 | 0 | 0 | 0 | 0 |
|  | Egg | 0 | **18.8 ± 32.5** | 0 | 0 | 0 | 0 | 0 | 0 | 0 | 0 |
|  | Euphausiids | 0 | 0 | 0 | 0 | 0 | 0 | 0 | 0 | 0 | 0 |
| | Other Copepodites | 0 | 0 | 0 | 1.5 ± 6.7 | 0.8 ± 3.7 | 0 | 0 | 0 | 0 | 0 |
| Flexion | Calanoid Copepodites | 0 | 0 | 1.9 ± 5.0 | 2.7 ± 8.6 | 1.5 ± 4.1 | 3.3 ± 16.9 | **3.1 ± 16.8** | 0.1 ± 0.4 | 0 | 0 |
| | Calanoid Nauplii | 1.0 ± 5.9 | **5.0 ± 7.3** | **9.3 ± 12.0** | **14.7 ± 26.0** | **2.2 ± 7.6** | 0 | 0 | 0 | 0 | 0 |
| | Cyclopoida Copepodites | 0 | 0 | 1.6 ± 4.1 | 1.0 ± 2.6 | 0.3 ± 1.1 | **2.3 ± 13.7** | 0 | 0 | 0 | 0 |
|  | Cyclopoida Nauplii | **7.3 ± 15.3** | **5.4 ± 11.3** | **19.2 ± 22.5** | 0 | 0 | 0 | 0 | 0 | 0 | 0 |
|  | Egg | 0 | **1.8 ± 6.5** | **3.9 ± 16.5** | 0 | 0 | 0 | 0 | 0 | 0 | 0 |
| | Euphausiids | 0 | 0 | 1.1 ± 6.4 | **1.4 ± 8.3** | **2.6 ± 15.4** | 0 | 0 | 0 | 0 | 0 |
| | Other Copepodites | 0 | 0 | **4.3 ± 17.7** | 0.5 ± 2.1 | 0 | 0 | **2.5 ± 14.7** | 0 | 0 | 0 |
| Postflexion | Calanoid Copepodites | 0 | 0 | 1.1 ± 3.0 | **2.5 ± 3.3** | **5.8 ± 11.3** | **6.9 ± 20.5** | **8.7 ± 24.7** | 0.3 ± 1.2 | **2.6 ± 9.3** | **1.4 ± 6.8** |
| | Calanoid Nauplii | 0 | 1.7 ± 3.2 | **10.3 ± 23.0** | **4.5 ± 18.2** | 0 | 0 | 0 | 0 | 0 | 0 |
| | Cyclopoida Copepodites | 0 | 0 | 0.2 ± 0.8 | 0.5 ± 1.7 | **4.4 ± 11.1** | **8.4 ± 20.6** | 0 | 0 | 0 | 0 |
| | Cyclopoida Nauplii | 0 | 1.1 ± 1.8 | **12.5 ± 21.1** | 0 | 0 | 0 | 0 | 0 | 0 | 0 |
| | Egg | **3.4 ± 16.5** | **4.6 ± 17.6** | **4.0 ± 15.1** | 0 | 0.1 ± 0.6 | 0 | 0 | 0 | 0 | 0 |
| | Euphausiids | 0 | 0 | 0 | 0 | 0 | **2.8 ± 9.8** | **2.1 ± 8.6** | 0.6 ± 2.3 | <0.1 ± 0.2 | 0.4 ± 2.1 |
| | Other Copepodites | 0 | 0 | **3.4 ± 16.6** | **2.1 ± 7.3** | 0.9 ± 3.0 | <0.1 ± 0.1 | 0.2 ± 1.2 | **2.4 ± 11.6** | 0 | 0 |

**Table 5:**
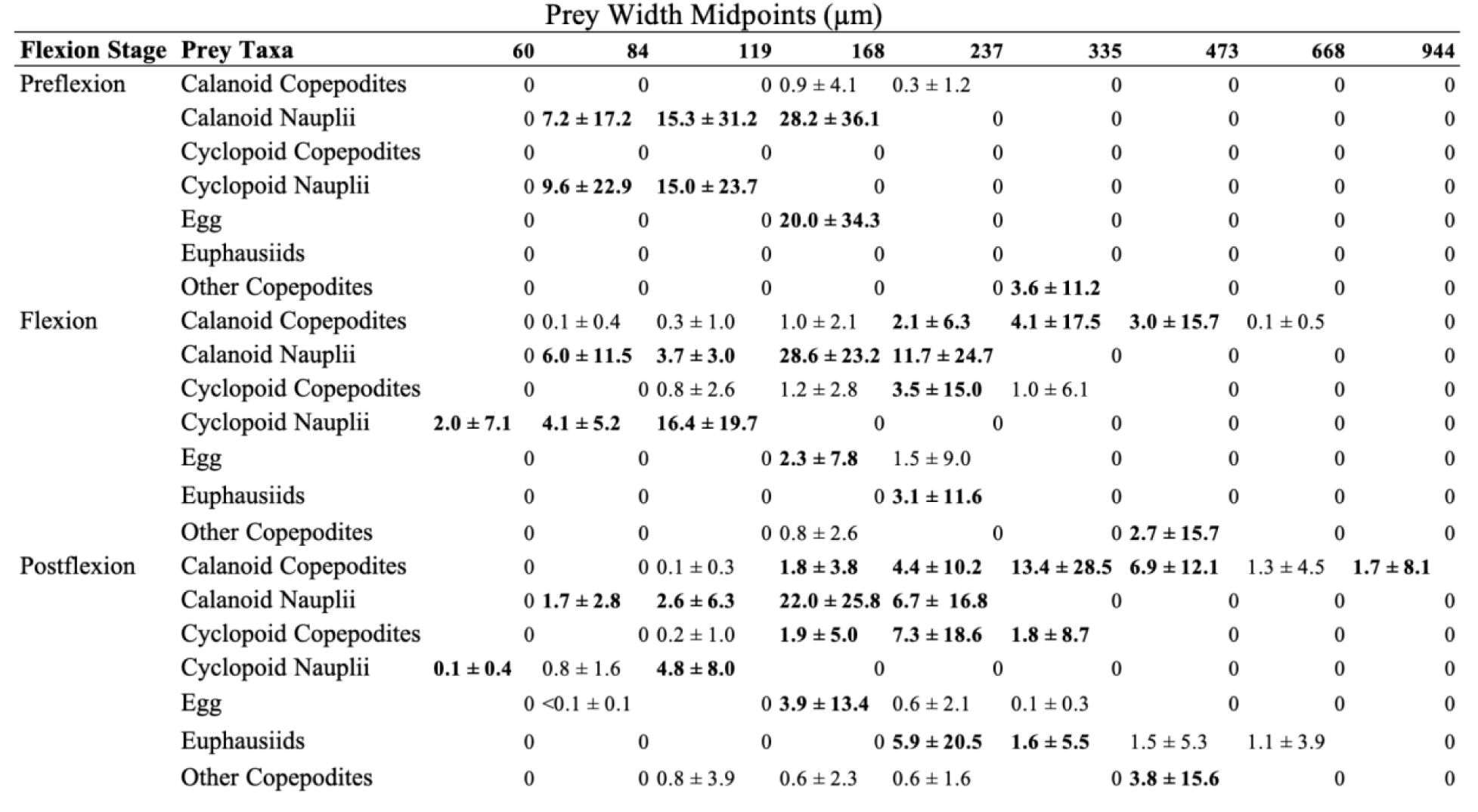
Chesson’s α-selectivity index calculations for nine logarithmic width classes with the seven most commonly consumed prey groups. Values are average α ± standard deviation. Bolded values represent taxa and size ranges that are positively selected for. Neutral preference is 1.59%.

### 3.4 Model Output

A positive relationship was observed between core radius and age (mean 0.47 ± 0.09, 89% posterior confidence interval (CI) 0.33-0.61) in model 1, although it did not follow a clear trend across stations (Figure 5). In the integrated diet and maternal investment hierarchical model 2 predicting standard length, age had the largest effect on standard length (mean 0.54 ± 0.08, 89% CI, 0.41-0.65; Figure 6). The next largest effect size was seen with core radius (mean 0.29 ± 0.05, 89% CI 0.12 - 0.28), closely followed by the relative contribution of Calanoid copepodite C biomass to the gut content (mean 0.18 ± 0.08, 89% CI 0.04-0.31). Note that due to the lack of additional predictor variables in model 1, the effect of core size on age is not directly comparable to its effect on standard length or recent growth in models 2 and 3. The proportion of Calanoid and Cyclopoid nauplii C biomass to the diet was found to have a negative effect on length (mean -0.13 ± 0.07, 89% CI -0.24- (-0.01)). The interaction between the proportion of nauplii biomass and age had a negative effect on standard length (mean -0.23 ± 0.07, 89% CI -0.35- (-0.11)), as did the interactive effect between the proportion of Calanoid copepodite biomass and age (mean -0.16 ± 0.07, 89% CI -0.28- (-0.04)). Effect sizes of random groupings did not follow a clear trend in the standard length model and there was a low strength of evidence for sampling location having a large effect. Relatively high uncertainty was also present in the random effect groups.

**Figure 5:**
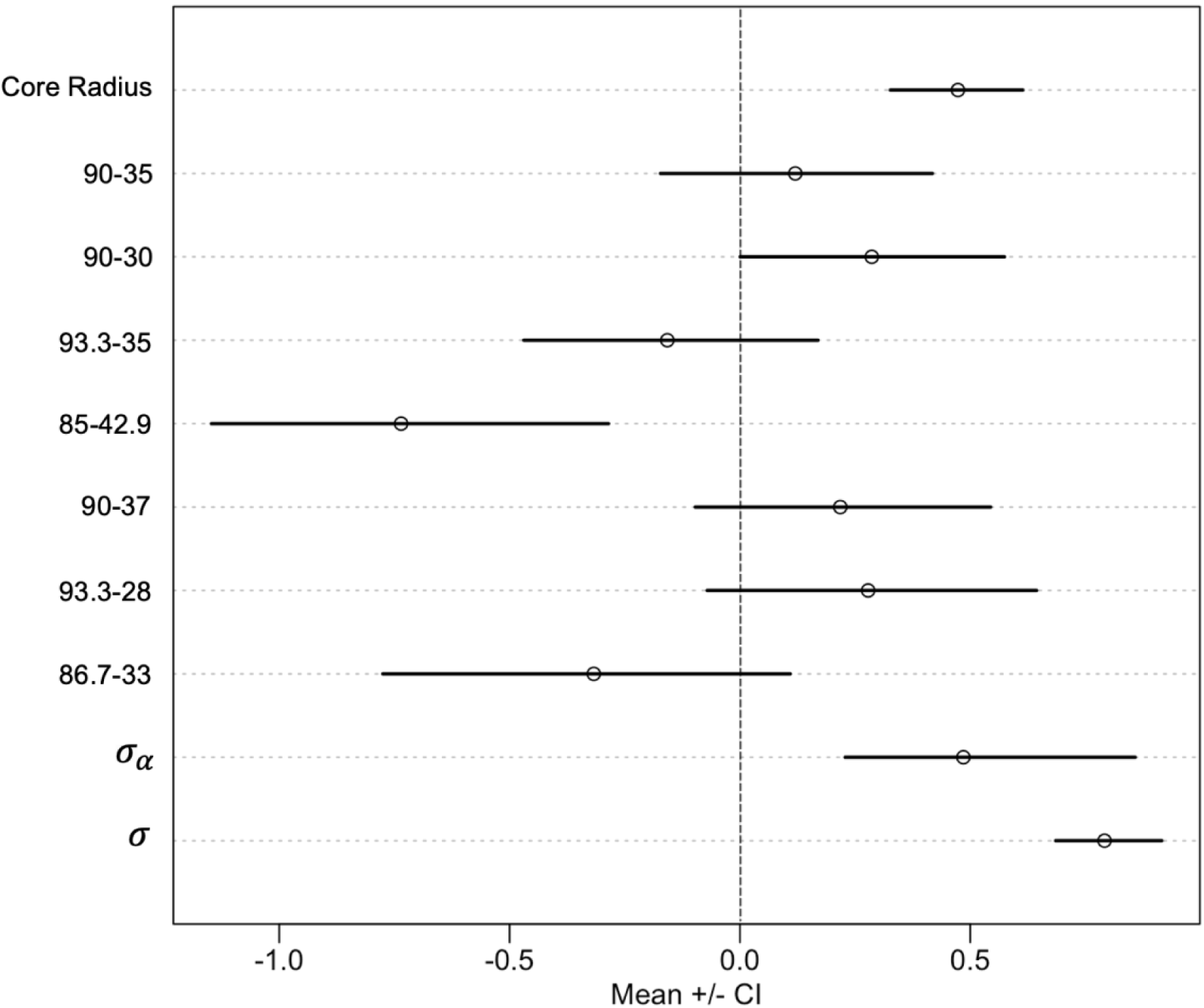
Coefficient plot of Bayesian hierarchical model 1 of the relationship between otolith core radius and age (days), describing mean effect sizes and 89% compatibility interval (CI). The sample stations are random effects, σ is standard deviation of normally distributed posterior distribution, and 𝜎_𝛼_ is standard deviation of the random intercept.

**Figure 6:**
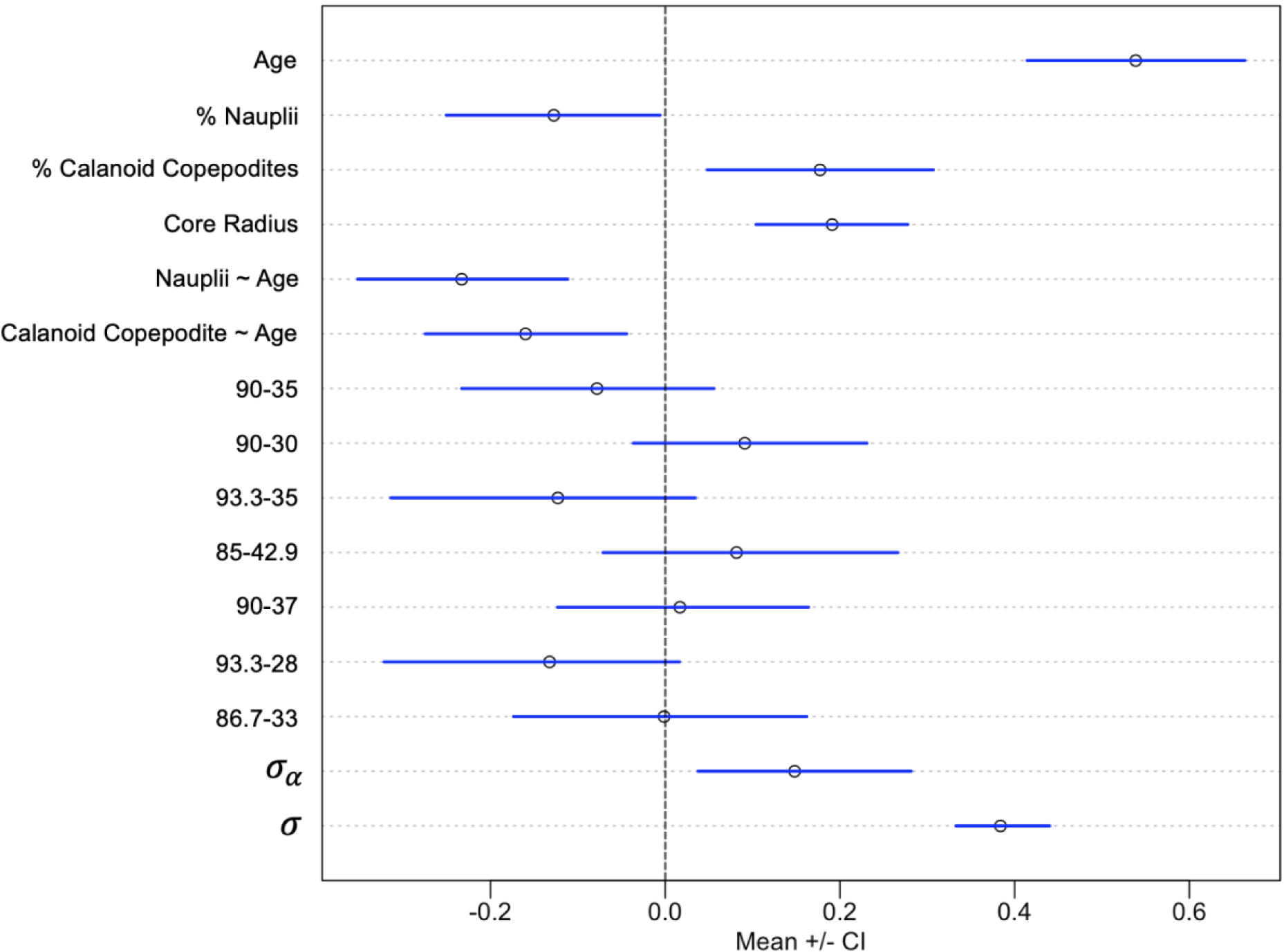
Coefficient plot of Bayesian hierarchical model 2 predicting standard length, describing mean effect sizes and 89% compatibility interval (CI). Age, % Nauplii, % Calanoid Copepodites, Core Radius, Nauplii∼Age, and Calanoid∼Age are fixed effects, with the last two representing the interactive effects of the respective diet parameters and age. The sample stations are random effects, σ is standard deviation of normally distributed posterior distribution, and 𝜎_𝛼_ is standard deviation of the random intercept.

For model 3 predicting recent growth, age was also the parameter most strongly correlated with recent growth of larvae (mean 0.42 ± 0.12, 89% CI 0.23-0.60; Figure 7). The next largest effect size was the proportion of Calanoid copepodite gut content C biomass (mean 0.31 ± 0.11, 89% CI 0.13-0.49), followed by core radius (mean 0.19 ± 0.08, 89% CI 0.06-0.31). The proportion of nauplii gut content C biomass had a slight negative correlation with recent growth (mean -0.06 ± 11, 89% CI -0.24-0.10). Negative effects were observed in the interactions of both nauplii C biomass and age (mean -0.13 ± 0.11, 89% CI -0.30-0.04) and Calanoid copepodite C biomass and age (mean -0.34 ± 0.11, 89% CI -0.52-(-0.16)). Effect sizes of random effects did not follow a clear trend in the recent growth model and had high uncertainty. Model fit was validated using posterior predictive checks, and models were observed to fit the data well (Supplementary Figure 1). Convergence diagnostics indicated good mixing of Markov chains.

**Figure 7:**
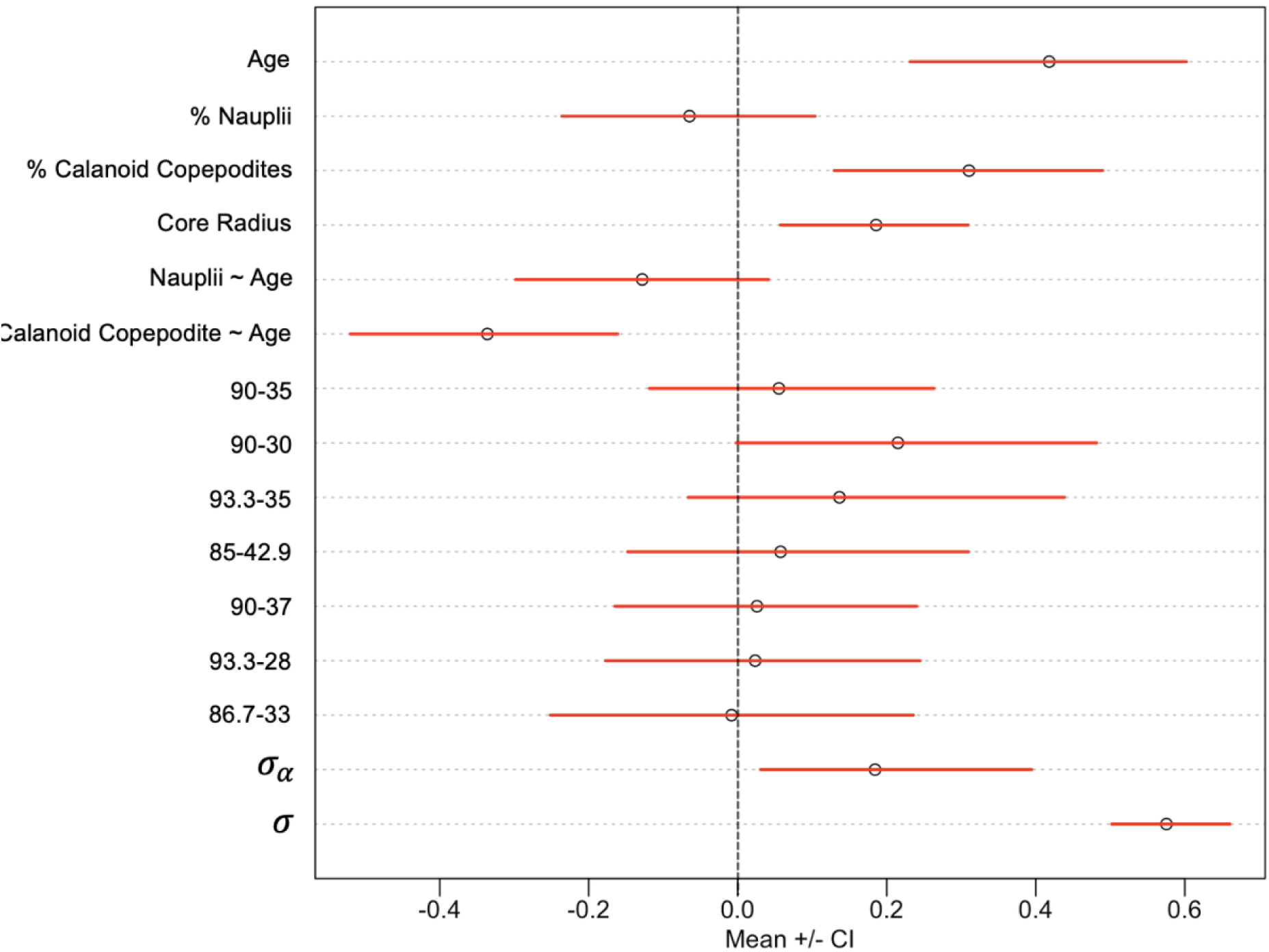
Coefficient plot of Bayesian hierarchical model 3 predicting recent growth, describing mean effect sizes and 89% compatibility interval (CI). Age, % Nauplii, % Calanoid Copepodites, Core Radius, Nauplii∼Age, and Calanoid∼Age are fixed effects, with the last two representing the interactive effects of the respective diet parameters and age. The sample stations are random effects, σ is standard deviation of normally distributed posterior distribution, and 𝜎_𝛼_is standard deviation of the random intercept.

Counterfactual plots were used to visualize the age-dependent effects of the proportions of different prey types on standard length and recent growth. In the standard length model (Figure 8), at an age of (-)1.96 (mean standardized age - 2SD) representative of younger larvae in the subsample, the relationship between increased Calanoid and Cyclopoid nauplii consumption and standard length was positive. At an age of 1.96 (mean standard age + 2SD) representative of older larvae in the subsample, the relationship between Calanoid and Cyclopoid nauplii consumption and length was negative. When examining the counterfactual effects of Calanoid copepodite consumption on standard length, the proportion of this prey parameter to the diet was positively correlated with length in younger larvae. A less distinct negative relationship was present with older larvae. When examining the counterfactual effects of these prey parameters on recent growth (Figure 9), a similar trend was apparent in that consumption of both prey groups tended to have positive correlations with recent growth in younger larvae and negative correlations in older larvae. However, the age-dependent relationships between nauplii biomass and recent growth were far less pronounced than between nauplii and standard length. The opposite was the case for Calanoid copepodite consumption, which had more pronounced positive effects on growth in younger larvae and more pronounced negative effects on older larvae.

**Figure 8:**
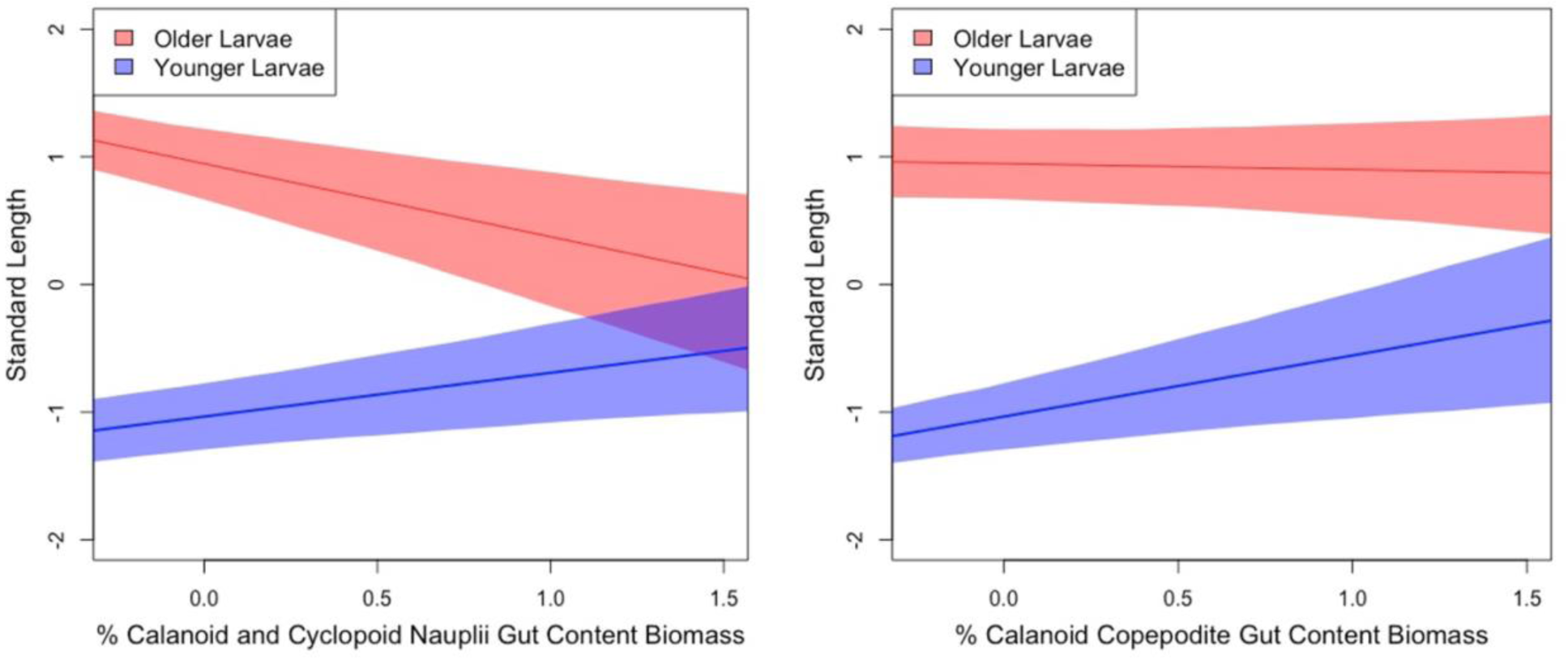
Counterfactual plots describing the relationship between the standardized proportion of Calanoid and Cyclopoid nauplii gut content biomass to the diet (left) and the standardized proportion of Calanoid copepodite gut content biomass to the diet (right) with standardized standard length in younger larvae (age = mean - 2 Standard deviations (SD)) and older larvae (age = mean + 2SD). Shaded regions represent 89% posterior intervals.

**Figure 9:**
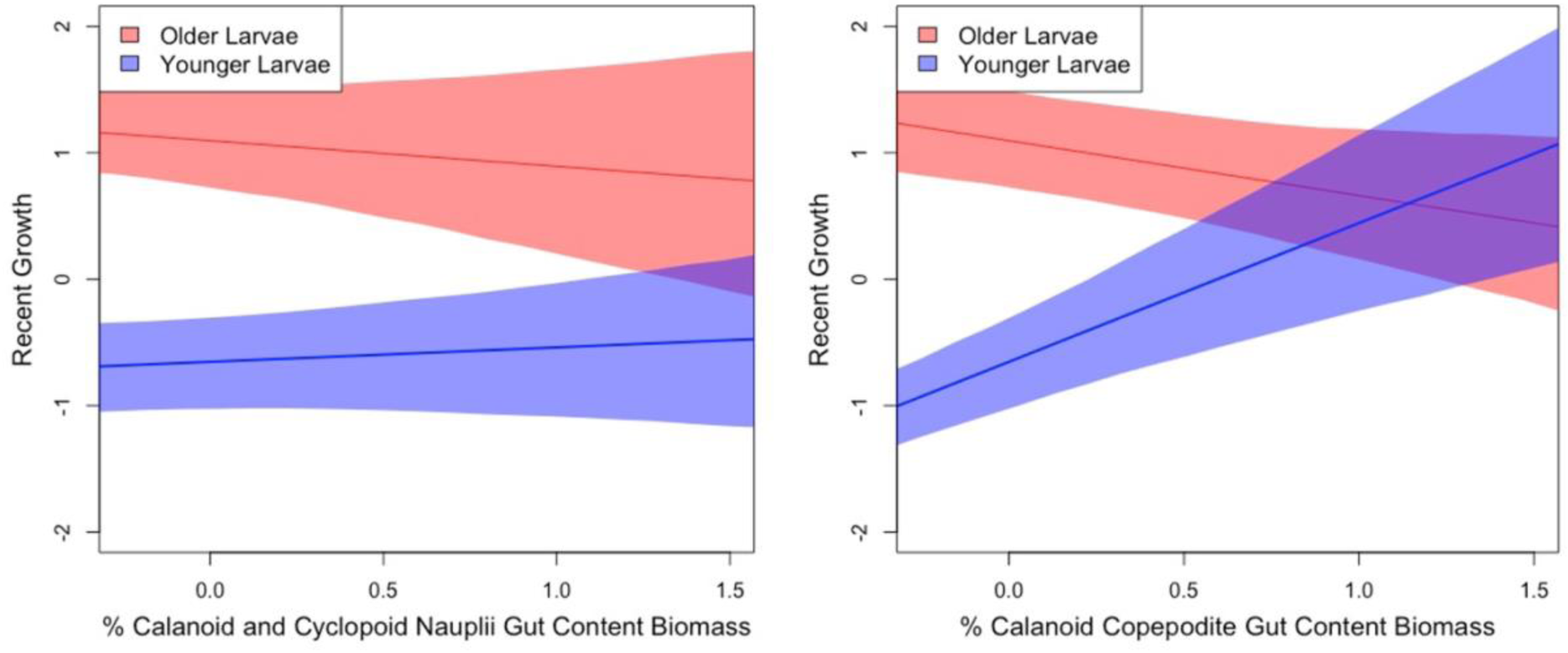
Counterfactual plots describing the relationship between the standardized proportion of Calanoid and Cyclopoid nauplii gut content biomass to the diet (left) and the standardized proportion of Calanoid copepodite gut content biomass to the diet (right) with standardized recent growth in younger larvae (age = mean - 2 Standard deviations (SD)) and older larvae (age = mean + 2SD). Shaded regions represent 89% posterior intervals.

## 4. DISCUSSION

Larval growth is governed by a confluence of factors that interact to shape the likelihood of an individual successfully recruiting to the adult population. Variability in growth has been long hypothesized to influence larval survival and fluctuations in year-class strength, with most studies finding slower-growing larvae having higher mortality (Chambers & Leggett 1987, Houde 1987, Miller et al. 1988, Hare & Cowen 1997, Houde 2008, Robert et al. 2023). While many studies have previously documented the diet and prey selection of larval fishes, the degree to which larval condition is shaped by the consumption of preferred prey has not been well studied (Robert et al. 2014). Thus, improving knowledge of how larval feeding influences growth in wild fishes at multiple life stages is essential for gaining a more mechanistic understanding of the ecological drivers of the “critical period” hypothesized over a century ago (Hjort 1914). In similar fashion, while maternal investment has emerged as a common driver of variation in size-at-birth and size-at-age in larval fishes as diverse as sardine, anchovy and rockfish (Garrido et al. 2015, Fennie et al. 2023, Robidas et al. 2022), studies investigating the joint effects of *in situ* size-at-birth and feeding ecology on larval ontogeny and condition are, to the best of our knowledge, non-existent. In this study, larval rockfishes shifted diet and selection towards specific prey taxa with ontogeny and exhibited active selection in response to prey availability in the surrounding environment. Both size-at-birth (otolith core size) and the consumption of specific prey had positive effects on larval size and recent growth. Preferred prey consumption affected size and recent growth differently at different life stages. Here, we discuss these results in the context of zooplankton ecology, regional oceanography, and previous early life history studies, with particular attention given to the challenges associated with interpreting causal mechanisms responsible for governing diet, size and growth in the larval stage.

### 4.1 Ontogenetic Shifts in Diet and Feeding Selectivity

Larval rockfishes shifted their diet with ontogeny, differing in the size of prey, number of prey, C biomass of prey, and type of prey consumed between growth stages. These results are consistent with previous studies that observed strong ontogenetic shifts in larval fish feeding (Murphy et al. 2012, Shiroza et al. 2021, Llopiz 2013). While feeding selectivity also changed throughout ontogeny, the most pronounced changes were between flexion and postflexion larvae. Changes in selection may mainly occur late in larval ontogeny, but it remains uncertain if larval fishes truly become more selective as they grow or if individuals that feed selectively early in ontogeny are more likely to survive to late larval stages. While older, larger larvae have better swimming ability than younger larvae and subsequently have better foraging capabilities (Margulies 1989, Majoris et al. 2019), variability in survival during the larval stage has also been attributed to selective mortality governed by phenotypic traits important for survival (Johnson et al. 2014). Prey selection early in ontogeny may dictate the proportion of larvae that survive to late larval stages, with observed feeding niche and selectivity trends in older larvae potentially being driven by the survival of individuals that had exhibited stronger selection for certain prey taxa earlier in life. However, due to insufficient data in this study to evaluate this hypothesis, we make the assumption that larval selection in our sample is plastic and older larvae experience greater changes in selection. Ontogenetic shifts in prey type remained within a narrower range of widths than lengths, indicating that prey width may be a greater limiting factor in selection than length. Selection towards developmental stages of copepod taxa remained within similar width ranges across ontogeny. This may be a function of esophagus width, which was not analyzed in this study but may influence size ranges of consumed prey and play a greater role in limiting consumption than mouth gape (Busch 1996). Size ranges of prey in guts might also be reflective of *in situ* abundances of different prey size ranges, with larvae selecting for size ranges found in the highest quantities in their environment. The lack of significant interspecific differences in diet and selectivity suggests that there is high overlap in diet between different species, and that niche partitioning may not occur until later in life. However, the low sample size of many species limits the conclusions that can be drawn.

### 4.2 Prey Availability Regulates Feeding Selectivity

Larval rockfish demonstrated strong selection for various life stages of copepods, particularly Calanoids. Calanoids of various life stages have been documented as a major food source of some *Sebastes* species (Sumida & Moser 1984, Nagasawa & Domon 1997, Swalethorp et al. 2015), but evidence of selection for this prey type within *Sebastes* has been limited to studies of Atlantic redfish (Anderson 1994). Changes in selection in response to changes in Calanoid availability have been reported for *Sebastes* in the North Atlantic, with poor larval condition and delayed metamorphosis associated with decreased consumption of Calanoids and increased consumption of *Oithona* spp. Cyclopoids (Anderson 1994). In our study, larval rockfish were observed to generally select more strongly for Calanoids than other copepod taxa such as Cyclopoids, including in sample stations in which Calanoids were not the most abundant copepod. Larvae were also observed to switch to alternative prey when Calanoids were locally scarce. Postflexion larvae selected strongly for Euphausiids and copepod eggs in stations which had relatively low abundances of Calanoid copepods (stations 93.3-28 and 93.3-35; Table 3).

Significant variation in selection between prey groups across sample stations indicates active selection by larval rockfishes, in which preference modulates in response to shifting prey group proportions in the surrounding environment (Shiroza et al. 2021). If selection were passive, it would have remained constant across sample stations irrespective of plankton community composition and spatial variability in the relative abundances of different prey taxa. While the benefits of active selection remain unclear, it is likely a function of some energetically favorable outcome associated with pursuing a particular prey type. Prey abundance facilitates increased predator encounter rates, and Calanoids are highly abundant in temperate ocean regions worldwide (Kozak et al. 2014). Historical species compositions of Calanoid populations have remained relatively stable in the CCE (Rebstock, 2001), but spatio-temporal changes in abundance, zooplankton community composition and larval distribution may nevertheless impact larval survival (Swalethorp et al., 2023, Cushing 1969).

Potential mechanisms for other prey taxa being selected differently than Calanoid nauplii and Copepodites also draw upon zooplankton physiology, ecology, and behavior. Euphausiids became selected for in late developmental stages in some sample stations and are known prey of early juveniles and even adults of some *Sebastes* (Chess et al. 1988, Bosley et al. 2014), but were largely avoided by early larvae and preyed upon in low quantities by older larvae. While more energy rich than copepods and potentially a good food source (Gorsky et al. 2010), Euphausiids generally fall outside of the size ranges most commonly selected for by rockfish larvae across ontogeny and are likely more difficult to acquire and handle. Cyclopoid copepodites were selected against on a sample population level, despite Cyclopoid nauplii being a preferred prey type. The preference of Calanoids over other copepodites may involve anatomical and behavioral characteristics that may make the latter more difficult to handle as prey. Cyclopoid copepods are primarily raptorial feeders that experience long periods of inactivity between strikes when compared to the highly motile suspension feeding often exhibited by Calanoids (Paffenhöfer et al. 1982, Williams & Muxagata 2006). Studies have suggested that larval strike rates may be triggered by more motile prey, potentially increasing encounter rates and facilitating increased visibility in turbid conditions (Sullivan et al. 1983, Buskey et al. 1993). Harpacticoids were largely absent from the gut contents despite being among the largest contributors to the *in situ* “Other Copepodites” category, and not only have long urosome spines that may deter predators but are characteristically particle-associated, likely making them more difficult for larval fishes to detect (Koski et al. 2005). While these characteristics of different taxa may be partially responsible for certain taxa being preferred over others, more research is needed to solidify understanding of the explanatory mechanisms driving feeding and selectivity in larval fishes.

Nevertheless, these findings highlight the importance of Calanoid copepods to larval rockfish and stress the importance of monitoring larval diets as a mechanism for improving understanding of the factors that contribute to interannual recruitment variability.

### 4.3 Ecological and Maternal Drivers of Condition and Growth

Analysis of diet and otolith core size yielded results that may increase understanding of the factors that influence variability in larval size and growth. Different prey items were associated with different condition responses. Calanoid copepodite gut content C biomass consistently displayed positive effects in the standard length and recent growth models, suggesting that Calanoid copepodite consumption may drive relationships between larval rockfish diet and condition. It also emphasizes the importance of considering taxonomic prey preferences when predicting the effects of diet on size, growth, and subsequently recruitment (Mayer & Wahl 1997, Castonguay 2008, Malca et al. 2022).

The positive correlations between otolith core size (a proxy for larval size-at-hatch/birth; Garrido et al. 2015, Fennie et al. 2023), and age, length and recent growth suggest that size-at-birth increases survivorship, as individuals born larger are more likely to survive to older ages, reach larger sizes, and grow faster. These results are consistent with trends observed in recent studies, in which core size was positively correlated with age, size, and condition in fishes with very different adult life history traits (European Sardine - Garrido et al. 2015, rockfishes - Fennie et al. 2023, Northern Anchovy - Robidas et al. 2022). More observational research needs to be conducted to discern the reason that larger larvae are more likely to survive. However, larger larvae may be more effective foragers by overcoming hydrodynamic feeding constraints imposed by low Reynolds’ Numbers (China & Holzman 2014). Hatching at larger sizes may thus provide a selective advantage throughout the larval stage by increasing prey capture efficiency.

Larval size-at-birth is most likely a function of maternal investment. Variability in maternal investment is generally attributed to parental condition, and numerous studies show that older, larger females produce better quality offspring (Berkeley et al. 2004, Sogard et al. 2008, Rodgveller et al. 2012). In the California Current Ecosystem, rockfish larvae otolith core size was recently observed to be influenced by oceanographic conditions experienced by the parents during parturition. When the parents were exposed to oxygen and nutrient-rich Pacific subarctic upper water (PSUW, i.e., California Current water), size-at-birth was larger (Fennie et al. 2023). The reason that parental condition is affected positively by PSUW is currently unknown, but there are multiple potential and non-exclusive reasons. For example, PSUW supports high primary and secondary production in the CCE, which may facilitate improved feeding conditions and higher quality prey for parents and larvae (Chelton et al. 1982, Miller et al. 2017). In addition, exposure to high oxygen PSUW has been hypothesized to improve the physiological condition of parents and enhance energy dedicated to producing offspring (Schroeder et al. 2019, Fennie et al. 2023). Improving our understanding of the relationship between size-at-birth, otolith core size and maternal investment in rockfishes, and the factors regulating maternal investment continues to be a promising route for elucidating mechanisms that affect fish recruitment (e.g., Hixon et al. 2014).

While initial larval size is unaffected by events that occur post-birth, reconciling causal pathways between diet and size is more challenging. Diet affects size by facilitating growth, but larger fish are also able to consume more food (Pirhonen et al. 2019). We attempted to determine the degree to which expected increases in food intake by older fish accounted for observed responses by using the proportions of important prey groups over the total amount of prey consumed by each larva. The positive correlations between the proportion of the total diet composed by Calanoid copepodites and both larval length and recent growth suggested that these trends were not simply generated by larger fish consuming disproportionately more food than smaller fish, but by consuming a greater proportion of a certain prey type relative to their total diet. We also included interactions between each diet parameter and age in the standard length and recent growth models, which generally showed stronger correlations between consumption of both preferred prey types and condition in younger larvae than in older larvae. This was particularly pronounced in younger larvae whose diets consisted of high proportions of copepodites. This supports the notion that increased consumption of Calanoid copepodites earlier in life may be advantageous for larval survival, particularly as it improves larval growth, while continued reliance on nauplii later in life may not be. Although Calanoid copepodites were most strongly selected for in the postflexion stage, they may provide a more energy-rich food source for larvae that are able to acquire and handle them than small nauplii (Rønnestad et al. 2013, Leech et al. 2021). The strength of evidence suggesting that some prey items were better predictors of size and growth than others followed a trend similar to that recently observed in young Atlantic redfish (Burns et al. 2021), in which consumption of the dominant and preferred prey, copepod eggs, had a weaker relationship with recent growth than more developed prey.

Our research indicates that larval condition is impacted by consumption of specific prey.

Although size at a given time is unlikely to be the result of growth induced by gut content biomass at that same time, gut content biomass in larval fishes likely reflects a larva’s foraging capacity over a span of days or even weeks. This notion is strengthened by the known sensitivity of larval fishes to food-poor conditions (Rice et al. 1987, Shan et al. 2008), with rapid starvation occurring in a few days or less without food (Hunter & Coyne 1982, Yin & Blaxter 1987, Garrido et al. 2015). Larvae are part of the broader plankton community and if hatched in a favorable feeding environment have the potential to stay in it with ontogenetic changes in feeding synchronous with prey development (Govoni et al. 2010, Gerard et al. 2022). Larvae were once thought to be passive particles with limited capacity to swim against ocean currents (Scheltema 1971), but recent studies have demonstrated the ability of larvae to exhibit some degree of active dispersal and directional swimming (Paris & Cowen 2004, Fisher et al. 2005, Leis 2006, Leis 2007) and thus may have the capacity to seek out patches of appropriate prey.

### 4.4 Conclusion and Future Directions

Prey type and the effects of otolith core size, a proxy for size-at-birth, impacted the condition of larval rockfishes in the Southern California Bight. Larvae selectively preyed on copepod nauplii and Calanoid copepodites, with increased consumption of Calanoid copepodites more strongly and positively correlated with size and growth than consumption of nauplii, particularly in young larvae. Otolith core size was also positively correlated with age, size and growth. Taken together, our study highlights the importance of larval size and feeding on specific preferred prey early in life with implications for larval survival and likely recruitment.

Future studies should center around solidifying causal relationships between diet and condition metrics in wild caught larvae to overcome challenges in interpreting how diet affects condition, identifying species-specific trends in feeding with larger sample sizes of different species, and improving understanding of why certain prey taxa are selected over others.

Examining how other indices of maternal condition in adult fishes relate to the otolith core sizes of offspring will aid in evaluating the degree to which core size can be reliably used as an index of maternal investment. In addition, future work should also aim to elucidate how ecological drivers of diet and maternal provisioning change in response to oceanographic regime shifts - particularly with regard to how larval success is affected by a warming ocean and how recent patterns of recruitment have confounded past understanding of how species fare in response to such changes (Jacox et al. 2016, Thompson et al. 2022). Lastly, the use of genetic techniques such as metabarcoding and environmental DNA bears great potential to not only increase the taxonomic resolution of preferred prey taxa but overcome some of the limitations of conventional stomach content analysis (Aguilar et al. 2017, Snider et al. 2022, Satterthwaite et al. 2023). Utilization of novel methods to improve sampling strength and confidence of prey identification will help solidify relationships between prey abundance and larval condition by accurately honing in on preferred species, improving the understanding of how diet and selectivity influence recruitment success.

## ACKNOWLEDGEMENTS

This work is funded by a NSF-RAPID Award (OCE-2053719). The authors would like to thank the captains and crew of the R/V Bob and Betty Beyster and R/V Shearwater, the NOAA Southwest Fisheries Science Center, Connor Coscino, Mika Carl, Celest Sorentino, Kelly Bishop, Dr. Noah Ben-Aderet, Dr. Mark Ohman, Dr. Noelle Bowlin, Dr. William Watson, Lauren Kittel-Porter, and Kamran Walsh’s MS thesis committee members Dr. Moira Décima and Dr. Colleen Petrik for their invaluable work and assistance on this project.

