## Supplemental material for "Diet and Size-at-Birth Affect Larval Rockfish Condition and Survival"

SUPPLEMENTARY MATERIAL

### Ambient Plankton Communities

Stations on the same transect lines and sampled in the same season (Figure 1) tended to be more similar in abundance contributions of different zooplankton (Supplementary Table 4) than stations sampled on different transects (mean dissimilarity between lines = 21.46%) and in different seasons (mean dissimilarity between seasons = 20.75%). Plankton community biomass trends were consistent in that larger dissimilarity was observed between stations on different transects (Supplementary Table 5) (mean dissimilarity between transects = 30.08%). However, different stations sampled during the same season were more dissimilar (mean dissimilarity = 25.71%) in prey taxa biomass contributions than different stations sampled on different seasons. Station 93.3-28, the only station sampled for *in situ* plankton community composition during both Winter and Spring, had a relatively low average dissimilarity between seasons in both abundance (16.48%) and biomass (14.33%).

The most abundant taxa in all stations were diatoms and protists, also accounting for the highest biomass in two of the five stations. Calanoid copepodites and nauplii were more abundant than Cyclopoids on transect line 90 and in station 93.3-28 (Winter), with the opposite true in the remaining stations. Euphausiids accounted for the highest carbon biomass per organism. Contributions of Poecilostomatoids (e.g. *Corycaeus* spp.), Harpacticoids (e.g. *Microsetella norvegicus*), copepods unidentified to a higher taxonomic resolution, and Cladocerans to total community abundance and biomass were relatively minor. Other zooplankton taxa were present in the original samples but excluded from here due to absences in the larval gut contents.

**Otolith Microstructure**

Otoliths were extracted from 19 preflexion, 30 flexion, and 31 postflexion larvae. Larvae that were younger than 3 days old and larvae with empty guts were excluded from subsequent analyses. The average age was 17.5 days post-extrusion ± 10.5 SD and the age range was 4-49 days post-extrusion. The average core radius was 15.7µm ± 2.7, with a range of 10.2 – 22.6 µm. The mean of the averaged three outer bands' widths was 2.63µm ± 1.4, with a range of 0.8 – 6.1 µm.

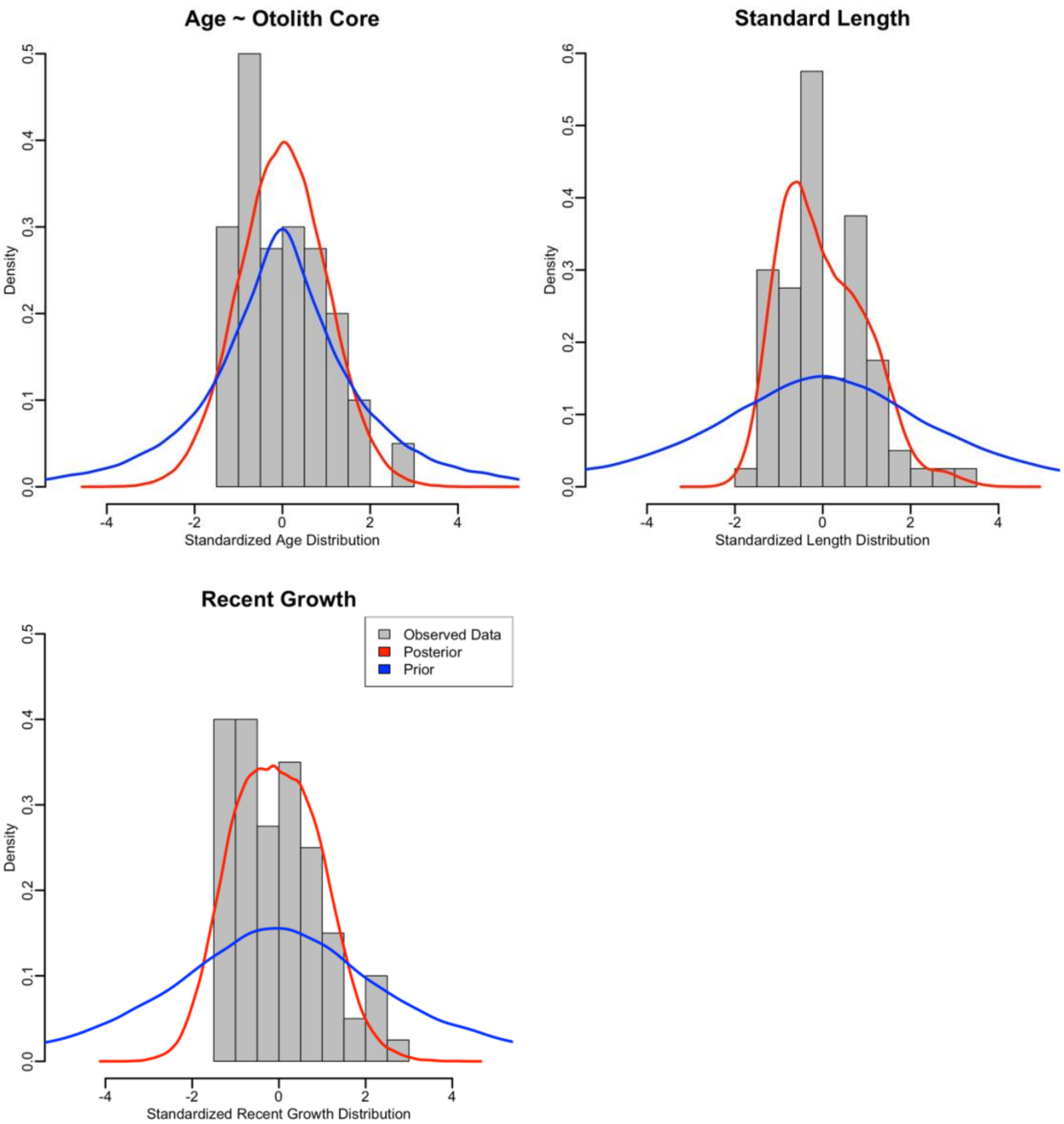


*Supplementary Figure 1: Posterior predictive checks for age (model 1; top left), standard length (model 2; top right) and recent growth (model 3; bottom) models displaying observed age, length and recent growth distributions overlaid with posterior (red) and prior (blue) distributions.*

*Supplementary Table 1: Length (L) to dry weight (DW) to carbon (C) conversion factors for larval rockfish prey taxa and in situ plankton community members.*

| Prey Group | Equation | C:DW | Reference |
|---|---|---|---|
| Calanoida Copepodite | $\ln(\mu g\ C) = 2.74 \times \ln(L_{\mu m})-16.41$ | 0.45 | Chisholm & Ruff 1990 |
| Cyclopoida, Other Copepodite | $\log_{10}(\mu g\ C) = 3.16 \times \log_{10}(L_{\mu m})-8.18$ | 0.45 | Hopcroft et al. 1998 |
| Poecilostomatoida Copepodite | $\ln(\mu g\ C) = 2.9 \times \ln(L_{\mu m})-17.5$ | | Satapoomin 1999 |
| Nauplii (all) | $\mu g\ C = 5.16 \times (L_{mm})^{2.372}$ | | Tanskanen 1994 |
| Diatoms and Protists | | | |
| Diatoms | $\mu g\ C = 1 \times 10^{-6} \times 0.288 \times BV_{\mu m3}{}^{0.811}$ | | Menden-Deuer & Lessard 2020 |
| Dinoflagellates | $\mu g\ C = 1 \times 10^{-6} \times 0.216 \times BV_{\mu m3}{}^{0.939}$ | | Menden-Deuer & Lessard 2020 |
| Other protists | $\log_{10}(\mu g\ C) = (-0.639+0.984 \times \log_{10}\pi(((L_{\mu m}/4)^2) \times (L_{\mu m}/3))))/(1 \times 10^6)$ | | Putt & Stoecker 1989 |
| Cladocerans | $\log_{10}(\mu g\ C) = 2.88 \times \log_{10}(L_{\mu m})-8.0767$ | | Shiroza et al. 2021 |
| Euphausiids (post-naupliar) | $\mu g\ C = 19.032 \times (L_{mm})^{1.484}$ | | Gorsky et al. 2010 |
| Egg | $\mu g\ C = 1.4 \times 10^{-7} \times BV_{\mu m3}$ | | Kiørboe et al. 1985 |

*Supplementary Table 2: Scientific and common names of larval rockfishes (Sebastes spp.) analyzed for gut content, proportion of the sample represented by each species, and proportion of stations where each species was present.*

| Species | Common Name | % of Total | % of Stations Present |
|---|---|---|---|
| *S. aurora* | Aurora | 0.621 | 14.286 |
| *S. chloristicus/rosenblatti* | Greenspotted/Greenblotched | 0.621 | 14.286 |
| *S. diploproa* | Splitnose | 0.621 | 14.286 |
| *S. ensifer* | Swordspine | 3.106 | 57.143 |
| *S. goodei* | Chilipepper | 0.621 | 28.571 |
| *S. helvomaculatus* | Rosethorn | 0.621 | 14.286 |
| *S. hopkinsi* | Squarespot | 11.801 | 85.714 |
| *S. jordani* | Shortbelly | 9.938 | 85.714 |
| *S. levis* | Cowcod | 1.242 | 14.286 |
| *S. macdonaldi* | Mexican | 1.242 | 28.571 |
| *S. melanostomus* | Blackgill | 2.484 | 42.857 |
| *S. miniatus* | Vermilion | 3.106 | 42.857 |
| *S. moseri* | Whitespeckled | 2.484 | 28.571 |
| *S. mystinus/entomelas* | Blue/Widow | 1.242 | 28.571 |
| *S. paucispinis* | Boccaccio | 1.242 | 28.571 |
| *S. rufinanus* | Dwarf-Red | 1.242 | 28.571 |
| *S. rufus* | Bank | 6.211 | 71.429 |
| *S. saxicola* | Stripetail | 1.863 | 42.857 |
| *S. semicinctus* | Halfbanded | 29.193 | 100.000 |
| *S. simulator* | Pinkrose | 9.938 | 100.000 |
| *S. wilsoni* | Pygmy | 2.484 | 42.857 |
| Unidentified *Sebastes* spp. | | 8.075 | 14.286 |

*Supplementary Table 3: spatial distributions of rockfish larvae growth stages selected for gut content analysis.*

| Station No. & Season | No. Preflexion | No. Flexion | No. Postflexion | Total per Station: |
|---|---|---|---|---|
| 90-35 Spring | 10 | 10 | 10 | **30** |
| 90-30 Spring & Winter | 10 | 10 | 19 | **39** |
| 86.7-33 Winter | 12 | 5 | 0 | **17** |
| 93.3-35 Spring | 7 | 5 | 2 | **14** |
| 93.3-28 Spring & Winter | 3 | 10 | 5 | **18** |
| 90-37 Spring | 3 | 10 | 7 | **20** |
| 85-42.9 Spring | 10 | 11 | 2 | **23** |
| **Total Per Growth Stage** | **55** | **61** | **45** | **161** |

*Supplementary Table 4: Spatial abundance distributions of zooplankton and phytoplankton prey field. Abundance is measured in counts per m3.*

| | Taxa Abundance Counts / m3 | | | | |
|---|---|---|---|---|---|
| **Taxa Group** | **93.3-28 Winter** | **90-35 Spring** | **90-37 Spring** | **93.3-28 Spring** | **93.3-35 Spring** |
| Calanoida Copepodites | 85.56 | 326.77 | 149.10 | 15.14 | 8.44 |
| Calanoida Nauplii | 149.16 | 123.82 | 24.01 | 23.89 | 18.56 |
| Cladocerans | 0.17 | 33.92 | 4.64 | 0.20 | 0.01 |
| Cyclopoida Copepodites | 55.26 | 20.02 | 10.88 | 22.86 | 12.91 |
| Cyclopoida Nauplii | 88.43 | 63.32 | 20.32 | 36.56 | 24.74 |
| Diatoms & Protists | 13792.45 | 1012.36 | 422.86 | 3044.32 | 1410.37 |
| Eggs | 19.21 | 17.66 | 12.88 | 9.62 | 14.24 |
| Euphausiid Nauplii | 1.13 | 1.18 | 1.28 | 0.00 | 0.00 |
| Euphausiids | 2.58 | 54.77 | 19.70 | 0.73 | 0.53 |
| Other Copepodites | 14.48 | 11.78 | 3.52 | 10.60 | 6.67 |
| Poecilostomatoida Copepodites | 1.04 | 33.97 | 13.68 | 1.28 | 1.93 |
| **Total Abundance m3** | **14209.46** | **1699.57** | **682.87** | **3165.21** | **1498.39** |

*Supplementary Table 5: Spatial biomass distributions of zooplankton and phytoplankton prey field. Biomass is measured in µg C per m3.*

| | Taxa Biomass µg C / m3 | | | | |
|---|---|---|---|---|---|
| **Taxa Group** | **93.3-28 Winter** | **90-35 Spring** | **90-37 Spring** | **93.3-28 Spring** | **93.3-35 Spring** |
| Calanoida Copepodites | 69.19 | 1766.53 | 367.63 | 22.77 | 11.28 |
| Calanoida Nauplii | 12.64 | 18.77 | 4.26 | 3.26 | 2.77 |
| Cladocerans | 0.12 | 36.00 | 4.23 | 0.11 | 0.00 |
| Cyclopoida Copepodites | 15.01 | 5.44 | 5.61 | 6.06 | 2.67 |
| Cyclopoida Nauplii | 7.56 | 5.12 | 1.45 | 2.83 | 1.98 |
| Diatoms & Protists | 52.13 | 42.66 | 14.37 | 75.27 | 87.62 |
| Eggs | 10.76 | 22.76 | 15.99 | 7.80 | 7.44 |
| Euphausiid Nauplii | 0.74 | 1.02 | 1.17 | 0.00 | 0.00 |
| Euphausiids | 107.13 | 2010.43 | 766.57 | 28.31 | 17.54 |
| Other Copepodites | 6.44 | 33.98 | 9.02 | 4.94 | 5.71 |
| Poecilostomatoida Copepodites | 1.33 | 77.03 | 36.38 | 1.65 | 4.65 |
| **Total Biomass m3** | **283.05** | **4019.77** | **1226.68** | **153.01** | **141.66** |
